# MUCOSECRETORY LUNG DISEASE: DIFFERENT ASSEMBLIES OF JAG1 AND JAG2 DETERMINE TRACHEOBRONCHIAL CELL FATE

**DOI:** 10.1101/2022.01.29.478334

**Authors:** Susan D. Reynolds, Cynthia L. Hill, Alfahdah Alsudayri, Scott W. Lallier, Saranga Wijeratne, ZhengHong Tan, Tendy Chiang, Estelle Cormet-Boyaka

**Author notes:** Contact Information: Susan D. Reynolds, PhD, Nationwide Children’s Hospital, 700 Children’s Drive, Columbus, OH 43205.

## Abstract

Mucosecretory lung disease compromises airway epithelial function and is characterized by goblet cell hyperplasia and ciliated cell hypoplasia. These cell types are derived from tracheobronchial stem/progenitor cells via a Notch dependent mechanism. Although specific arrays of Notch receptors regulate cell fate determination, the function of the ligands Jagged1 (JAG1) and Jagged2 (JAG2) is unclear. This study used primary human bronchial air-liquid-interface cultures, gamma secretase inhibition, and neutralizing antibodies to show: 1) JAG1 and JAG2 were necessary for secretory progenitor cell fate determination; 2) JAG2 suppressed squamous differentiation; and 3) pausing of the ciliated cell differentiation process after Notch inhibition. Histological, cell fractionation, cell surface biotinylation, and ubiquitination analyses demonstrated that all cells were JAG1 positive but that little JAG1 was present on the cell surface. In contrast, JAG2 was expressed in a positive-negative pattern and was abundant on the cell surface. Glycogen synthase kinase 3 (GSK3) and tankyrase inhibition studies showed that GSK3 regulated JAG2 trafficking, and that this mechanism was WNT-independent. Collectively, these data indicate that variation in JAG2 trafficking creates regions of high, medium, and low ligand expression. Thus, distinct assemblies of JAG1 and JAG2 may regulate Notch signal strength and determine the fate of tracheobronchial stem/progenitor cells.

Graphical Abstract
Different assemblies of JAG1 and JAG2 may determine Notch signal strength and cell fate within the tracheobronchial epithelium. A cell which interacts with JAG1+ cells (blue squares) receives a low Notch signal (light yellow square). A cell which interacts with a mixture of JAG1+ and JAG1+/JAG2+ cells (purple squares) receives a medium (med) Notch signal (medium yellow square). A cell which interacts with JAG1+/JAG2+ cells receives a high Notch signal (bright yellow square).

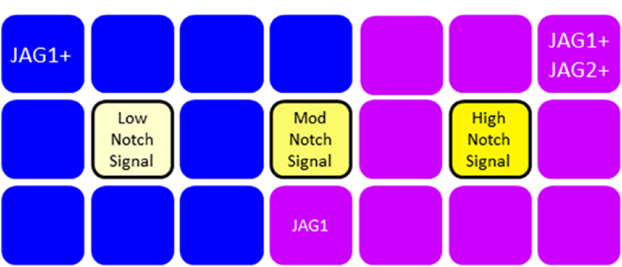

## INTRODUCTION

The conducting airway epithelium protects the lung from inspired environmental agents via mucociliary clearance. Effective removal of pathogens, irritants, and toxins (including cigarette smoke and air pollutants) requires the coordinated activity of two specialized airway epithelial cell types, the multiciliated cell (termed a ciliated cell) and secretory cells (including goblet cells and club cells). These cell types are descendants of the tracheobronchial epithelial tissue specific stem cell (TSC) which regenerates the pseudostratified airway epithelium and resides in the trachea and bronchi of the mouse and upper respiratory tract of humans. Lineage-tracing in mice (1–4) and clonal analysis in human (5) indicated that the TSC was a basal cell subtype.

Previous studies reported that tracheobronchial ciliated and secretory cells were present in approximately equal numbers and that their frequency was regulated by Notch signaling (reviewed in (6)). Specifically, the Notch Lateral Induction Pathway (Notch pathway) regulates basal and secretory cell differentiation (reviewed in (7)). Notch signaling occurs between adjacent cells which express Notch ligands (Jagged (JAG) 1 or 2 or Delta-like (DLL) 1, 3, or 4) or Notch receptors (NOTCH1, 2, 3, or 4) (6, 8). Following receptor ligation, ADAM10 or ADAM17 cleaves NOTCH at an extracellular site and the γ-secretase complex cleaves NOTCH at an intracellular site. This process releases the NOTCH intracellular domain which forms a transcriptional complex with several proteins including RBPJK and activates expression of the HES and HEY family of transcriptional repressors (9–11).

Chronic lung disease is frequently associated a disruption of mucociliary clearance. In asthma, chronic obstructive pulmonary disease (COPD), chronic bronchitis (CB), and idiopathic pulmonary fibrosis (IPF) goblet cell hyperplasia and excess mucus secretion drive the mucosecretory phenotype. Single cell RNAseq (scRNAseq) analysis of asthma, IPF, and COPD indicated that Notch signaling was aberrant (12–14) and functional studies indicated that cigarette smoke, an agent which increases risk of developing COPD, CB, or IPF, activated Notch signaling (15). Mucociliary clearance can also be disrupted by ozone and cigarette smoke exposure or infection with SARS-CoV2 and/or respiratory syncytial virus. These agents damage the ciliated cell, often resulting in cell death and ciliated cell hypoplasia. Notch pathway components were identified as susceptibility genes for lung damage (16), viral entry (17), and hyperinflammation (18). Consequently, treatments which specifically target Notch pathway components have the potential to normalize ciliated and goblet cell frequency, restore mucociliary clearance, and improve the health of people with chronic lung disease.

Several studies defined the Notch receptors responsible for generation of ciliated and secretory cells in mouse and human (13, 19–23). However, the Notch ligand(s) that is responsible for pathway activity is unclear. Studies in mice indicate that DLL ligands regulate generation of neuroepithelial cells but are unlikely to be involved in ciliated and secretory cell differentiation (24). In contrast, over-expression of *Jag2* mRNA relative to *Jag1* in basal cells (21), treatment-dependent increases in *Jag1* and *Jag2* mRNA (25), and co-localization of the two mRNAs (24, 26) suggested overlapping roles for JAG1 and JAG2 in basal cell fate determination. Conditional knockout of *Jag1* or *Jag2* early in mouse lung development supported this overlapping mechanism but other studies identified JAG2 as the dominant ligand (24, 26). Finally, a specific role for JAG1 in maintenance of secretory cell fate and suppression of ciliated cell fate was indicated by experiments involving *Jag1* overexpression in immortalized human basal cells (27), analysis of *Jag1* knockout in mice (28), and treatment of mice with a JAG1 neutralizing antibody (29) or a *Jag1* antisense oligonucleotide (30).

A potential explanation for the disparate Jag results is the focus on the fully differentiated ciliated or goblet cell. Ciliated cell subtypes have been identified by scRNAseq (31, 32), while histological analysis of the mouse tracheobronchial epithelium (33) and the Xenopus epidermis (reviewed in (34)) defined ciliated cell differentiation intermediates. These intermediates can be detected by immunostaining for acetylated tubulin (ACT, (35)). Similarly, multiple secretory cell subtypes have been identified, with the MUC5B-expressing goblet cell being the major subtype (36). Goblet cell differentiation intermediates can be identified by immunostaining for MUC5B.

This study examined Notch signaling during differentiation of tracheobronchial TSC and progenitor cells and focused on roles for JAG1 and JAG2. Cell fate determination was evaluated using human bronchial air-liquid-interface cultures, a well-described and widely-utilized model of the tracheobronchial epithelium (37). Treatment with γ-secretase inhibitors and analysis of differentiation intermediates was used to examine Notch regulation of cell fate. Roles for JAG1 and JAG2 in differentiation of TSC and the secretory progenitor were evaluated using histological and biochemical approaches. JAG1 and JAG2 necessity was examined using a JAG1 peptide and anti-JAG1 and anti-JAG2 neutralizing antibodies.

## METHODS

### Human

*TSC recovery, expansion, and ALI culture:* Human bronchial basal cells were recovered as previously described (38, 39). The modified conditional reprogramming culture (mCRC) method (39) was adapted from Suprynowicz et al (40). The major change from this protocol was the use of irradiated NIH3T3 fibroblast feeder layers. Human basal cells were differentiated using previously published methods (39). Initial studies used three culture media: Wu (41), Half & Half (H&H, (35), or complete Pneumacult (Stem Cell Technology, Vancouver, BC, Canada). Culture medium is defined in Results and in Figure Legends.

### ALI culture treatment protocol

The specific treatment and analysis days and concentration of agonists/antagonists are presented in Results and Figure Legends. Small molecule agonists and antagonists were purchased from Tocris (Minneapolis, MN) and dissolved in DMSO at 1000x the treatment concentration. Vehicle controls were treated with an equal volume of DMSO. Jag1 peptide was purchased from Anaspec (Freemont, CA) and dissolved in 1x Phosphate Buffered Saline (PBS) at 500x the treatment concentration. Vehicle controls were treated with an equal volume of PBS. Neutralizing antibodies specific for Jag1 (anti-JAG1.b70) and Jag2 (anti-JAG2.b33) were provided by Genentech (29) and were dissolved in PBS at 100x the treatment concentration. Vehicle controls were treated with an equivalent volume of PBS.

### Immunofluorescence analysis and quantification

Differentiation was evaluated in ALI cultures. Cultures were fix with 4% paraformaldehyde/1% sucrose/1x PBS. For analysis of intracellular markers cultures were permeabilized and blocked with 0.1% NP40/5% bovine serum albumin/1x PBS. For analysis of cell surface markers, cultures were fixed with 10% neutral buffered formalin and blocked with 5% bovine serum albumin/1x PBS. Goblet cells and differentiation intermediates were identified by MUC5B staining using rabbit-anti-MUC5B (Sigma, 1/500). Ciliated cells and differentiation intermediates were identified by ACT staining using mouse-anti-ACT (Sigma, 1/8000). JAG1 was evaluated using a mouse-anti-JAG1 C-terminal specific antibody (BD Biosciences #612346, 1/50). JAG2 was evaluated using a rabbit-anti-JAG2 N-terminal specific antibody (Cell Signaling C23D2, 1/50) and a rabbit-anti-JAG2 C-terminal specific antibody (Cell Signaling C83A8, 1/50). HES1 was evaluated using a goat-anti-HES1 (Santa Cruz #sc-13842, 1/40). Rbpjk was evaluated using a rabbit-anti-RBPJK (Sigma #AB5790, 1/100). Primary antibodies were detected with ALEXA-488 or ALEXA-594 labeled secondary antibodies (Jackson Immunological, 1/500). Nuclei were detected with DAPI. Cell type frequency was quantified as previously described (39).

### Cell fractionation

Cells were separated into cytoplasmic and nuclear fractions using a NE-PER kit (Thermo Scientific, P178833). 4x Laemmli’s Buffer/10% β-mercaptoethanol (BME) was diluted 1/4 into the protein fraction and the mixture was incubated at 96 °C for 10 minutes.

### Cell lysis

RIPA buffer (42) was used to lyse cells for immunoprecipitation, poly-ubiquitination, and western blot analysis. The apical and basal surfaces of ALI cultures were washed with Hams/F12 at 4 °C for 10 min. Wash solution was aspirated and 270 µl RIPA buffer was added to the apical surface. The cell layer was disrupted with a p200 pipette tip and then incubated for 20 min at 4 °C. The lysate was collected in a 1 ml microcentrifuge tube, sonicated 5 x 30 sec in a Bioruptor (Diagenode) at 4 °C, and centrifuged at 11,176g for 10 min at 4 °C. The supernatant was stored at -80 °C.

### Immunoprecipitation

Immunoprecipitation was performed using a modified version of the Pierce Protein Magnetic Bead protocol (Thermo 88848). For each sample, 125 µl of beads were pre-washed three times with RIPA Buffer and resuspended in RIPA Buffer with proteinase (Roche; 11836153001) and phosphatase inhibitors (Roche; 04906837001). Washed beads (25 µl) were added to 400-600 ug protein per sample and the lysate was cleared by rocking for 1 hr at 4 °C.

The cleared lysate was incubated with a titration-determined amount of antibody (usually 1:100). This mixture was rocked for 1 hr at 4 °C. The remaining 100 µl of washed beads were added to the sample and rocked for 2 hr at 4 °C. The beads were collected using a neodymium magnet, washed three times with RIPA Buffer, resuspended in 1x Laemmli’s Buffer plus 2.5% BME, and incubated at 96 °C for 10 minutes. The supernatant was kept as Unbound Fraction.

### Cell surface biotinylation

A modified version of the Pierce Cell Surface Protein Biotinylation and Isolation Kit (Thermo A44390) protocol was used to isolate surface proteins. The recommended biotinylation incubation time was increased to 45 minutes and incubations were done on ice.

### Poly-ubiquitination

Total Poly-ubiquitinated proteins were isolated using a LifeSensors Tandem Ubiquitin Binding Entities (TUBEs) Kit (LifeSensors, UM411M) and a modified protocol. The cell lysate was incubated with 100 µl washed, unconjugated beads at 4 °C on a rotating platform for 30 minutes. A bead titration study demonstrated an optimal ratio of 100 µl TUBE-conjugated beads per 1-2 mg protein in a total of 1 ml lysis buffer. The cleared lysate and washed TUBE beads were incubated at 4 °C on a rotating platform for 30 minutes. The TUBE beads were collected using a magnet, washed three times with PBST, and divided into two aliquots. One aliquot was treated with buffer and the second aliquot was treated with 2 µl of 10 µM broad- spectrum deubiquitinase for 2hr at 30°C with rocking. The beads were resuspended in 1x Laemmli’s Buffer/2.5% BME and incubated for 10 minutes.

### Western blot analysis and quantification

Protein concentration, gel electrophoresis and transfer, and blotting were done as previously reported (42). In some instances, blots were stripped by incubation with 1X Western Reprobe (G Biosciences) for 45 minutes at 55 °C followed by three washes with Tris buffered saline with TWEEN-20 (TBS-T, Cell Signaling Technology) at room temperature. Antibodies used for western blot studies: JAG2-N-terminal (Cell Signaling C23D2, 1:1000); JAG2-C-terminal (Cell Signaling C83A8, 1:1000); GSK3 (Millipore 05-412, 1:2000); pGSK3 (Millipore 0-413, 1:1000); 14-3-3 (Millipore AB9748-I, 1:2000); TOPO II (Abcam 109524, 1:10000); Sodium Potassium ATPase (NAK, Cell Signaling 3010S, 1:2500); JAG1-C-terminal (BD Biosciences #612346, 1:1000); α-Tubulin (Sigma T6793, 1:2000); N-terminal β-catenin Cell Signaling #9581X, 1:1000); C-terminal β-catenin (BD Transduction #610154, 1:2000); and Keratin 5 (BioLegend #905501, 1:2000).

### RNA preparation and RNAseq gene expression analysis

RNA was purified using a RNeasy Kit (Qiagen,74106) according to the manufacturer’s directions. RNA-sequencing (RNAseq) was conducted by the Institute for Genomic Medicine at Nationwide Children’s Hospital. On average, 32 million paired-end 151 bp RNA-Seq reads were generated for each sample (the range was 25 to 37 million). Each sample was aligned to the GRCh38.p9 assembly of the Human reference from NCBI (http://www.ncbi.nlm.nih.gov/assembly/GCF_000001405.35/) using version 2.6.0c of the RNA-Seq aligner STAR (http://bioinformatics.oxfordjournals.org/content/29/1/15). Transcript features were identified from the GFF file provided with the assembly from Gencode (v28) and raw coverage counts were calculated using featureCounts (http://bioinf.wehi.edu.au/featureCounts/). The raw RNA-Seq gene expression data was normalized and post-alignment statistical analyses were performed using DESeq2 (http://genomebiology.com/2014/15/12/550) and custom analysis scripts written in R. Comparisons of gene expression and associated statistical analysis were made between different conditions of interest using the normalized read counts. All fold change values are expressed as test condition / control condition, where values less than one are denoted as the negative of its inverse (note that there will be no fold change values between –1 and 1, and that the fold changes of “1” and “-1” represent the same value). Transcripts were considered significantly differentially expressed using a 10% false discovery rate (DESeq2 adjusted p value <= 0.1). GSEA was used to identify treatment dependent changes in signaling.

### scRNAseq data analysis

Single cell RNAseq methods and analytical techniques are presented in detail in (13) and (43). The raw reads were processed with Cell Ranger software 3.0.2 (10X Genomics) using hg38 transcriptome reference from GENCODE 25 annotation, and a digital expression matrix was generated with cell barcodes as rows and gene identities as columns. Each dataset was run with SoupX analysis package to remove contaminant ‘ambient’ RNA derived from lysed cells during isolation and capture (44). To minimize doublet contamination, for each dataset the Scrublet package was used to detect neotypic doublets (45) and a quantile thresholding was performed to identify high UMI using a fit model generated using multiplet’s rate to recovered cells proportion, as previously described (43). For all data, quality control and filtering were performed to remove cells with low number of expressed genes (threshold n>=200) and elevated expression of apoptotic transcripts (threshold mitochondrial genes < 15%). Only genes detected in at least 3 cells were included. Data were normalized using the Scran package (46), using a low resolution clustering to use as input for the size factor estimation and then log transformed. Data were processed with Principal Component Analysis (PCA) using the 5000 most variable genes as input, and clustering was performed with the Leiden algorithm. Batch correction for the two samples was performed with the package BBKNN (Polanski et al., 2019). Partition-based graph abstraction (PAGA) (47) was added to Uniform Manifold Approximation and Projection (UMAP) with the option init_pos=’paga’ (48) to obtain a two-dimensional visualization. To identify differentially expressed genes between clusters and representative of both datasets, Model-based Analysis of Single-cell Transcriptomics (MAST) (49) was used. To characterize cell subsets in our integrated dataset, previously published lung cell subtype specific gene lists (43) were used to create cell type-specific gene signatures using a strategy previously described (50). The steps described above were generated using Seurat 3.0 in R 4.0 or Scanpy 1.7.0 in Python 3.8.5. Plots were generated within Seurat and Scanpy or using the ggplot2 package.

### Statistics

All statistical analyses were performed using Graph Pad Prism. Data normality was evaluated by the Shapiro-Wilk test, the Anderson-Darling test, and/or the D’Agostino & Pearson test. For normally distributed data sets, differences were evaluated using Student’s t-test. For non-normally distributed data sets, differences were evaluated by the Mann-Whitney test and data are presented as the median and the interquartile range. Trends were analyzed by regression analysis. The definition of center and dispersion precision measures are indicated in the Figure Legends. Data sets containing multiple variables were analyzed by analysis of variance (ANOVA) and a post hoc Tukey test (normally distributed data sets) or Kruskal Willis test (non-normally distributed data sets). Sample size is indicated in the Figure Legends. P-values < 0.05 were considered significant.

### Study approval

The Institutional Review Board at Nationwide Children’s Hospital approved the human studies. Written informed consent and assent was obtained from every participant.

## RESULTS

### Ciliated and goblet cell differentiation

Since kinetics and cell type frequency vary across model systems, differentiation was evaluated in three differentiation media. Quantification demonstrated that cell density did not vary as a function of medium or time (SFig 1A) and that the frequency of ciliated cells increased significantly between differentiation days 7 and 14 in all three media (SFig 1B). The frequency of mature goblet cells did not vary between differentiation days 7 and 14 in Wu or complete Pneumacult medium but increased significantly in Half&Half (H&H) medium (SFig 1C). Based on the finding that H&H medium supported an increase in ciliated and goblet cell frequency over days 7-14 and generated similar numbers of goblet and ciliated cells on day 14, this medium was used to evaluate the signaling mechanisms that regulate production of ciliated and goblet cells.

### Intermediate ciliated and goblet cell phenotypes

Previous work demonstrated that ciliated cell differentiation involved a set of intermediate phenotypes (reviewed in (34)). In Stage I, cells exit the cell cycle and express a primary cilium (SFig 2A-B). This phenotype was detected at early time points and the frequency of cells with a primary cilium peaked on differentiation day 6 (Fig 1A). During Stages II and III, new centrioles are generated and dock at the apical cell surface. Cells that contained many basal bodies and occasional short cilia (bristle cells, SFig 2C-D) were identified on days 8-12 (Fig 1B). Finally, during Stage IV, each docked centriole (now termed a basal body) nucleates a motile 9+2 ciliary axoneme and the ciliary appendage is formed. Cells defined many long motile cilia (ciliated cells, SFig 2E-F) were detected on days 8-12 (Fig 1C).

**Figure 1:**
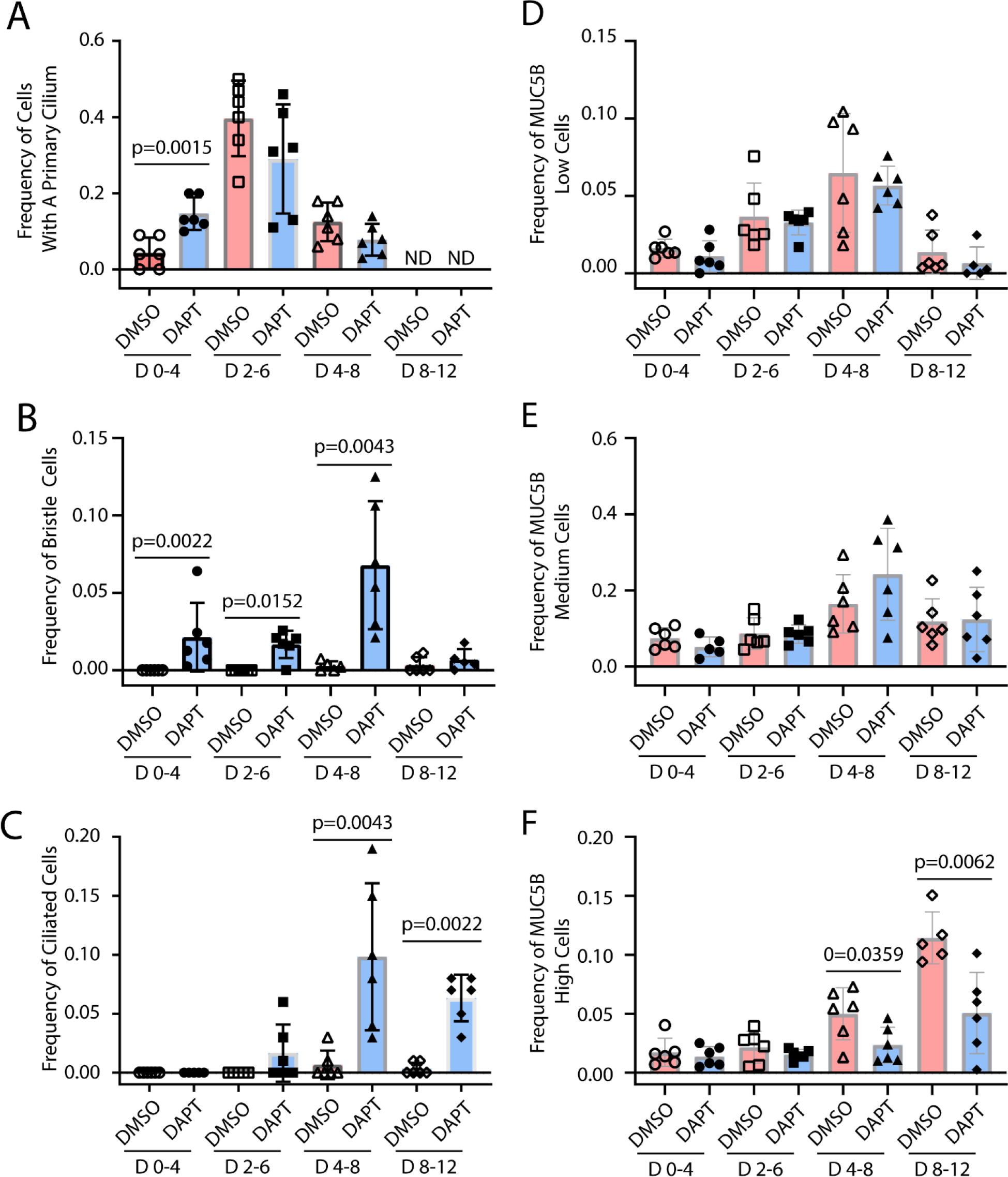
Notch regulation of ciliated and goblet cell differentiation. Human bronchial basal cells were differentiated in air-liquid-interface cultures using H&H medium. Cells were treated with vehicle (DMSO) or 25 µM DAPT as follows: treatment on differentiation days 0 and 2 and fix on day 4, treatment on differentiation days 2 and 4 and fix on day 6, treatment on differentiation days 4 and 6 and fix on day 8, or treatment on differentiation days 8 and 10 and fix on day 12. Dual immunofluorescence was used to detect acetylated tubulin and MUC5B. Nuclei were stained with DAPI. **A.** Frequency of cells with a primary cilium. **B.** Frequency of bristle cells. **C.** Frequency of ciliated cells. **D.** Frequency of MUC5B-low cells. **E.** Frequency of MUC5B-medium cells. **F.** Frequency of MUC5B-high cells. Mean ± SD, N=6.

Goblet cells also exhibited three morphological phenotypes that were defined by low, medium, and high expression of MUC5B (SFig 2G-H). All three intermediates were detected at all time points (Fig 1D-F). The frequency of MUC5B-low and MUC5B-medium cells peaked on day 8 while the frequency of MUC5B-high cells increased through day 12 (Fig 2F). These data indicated that the first wave of ciliated and goblet cell differentiation was completed the process by day 12.

**Figure 2:**
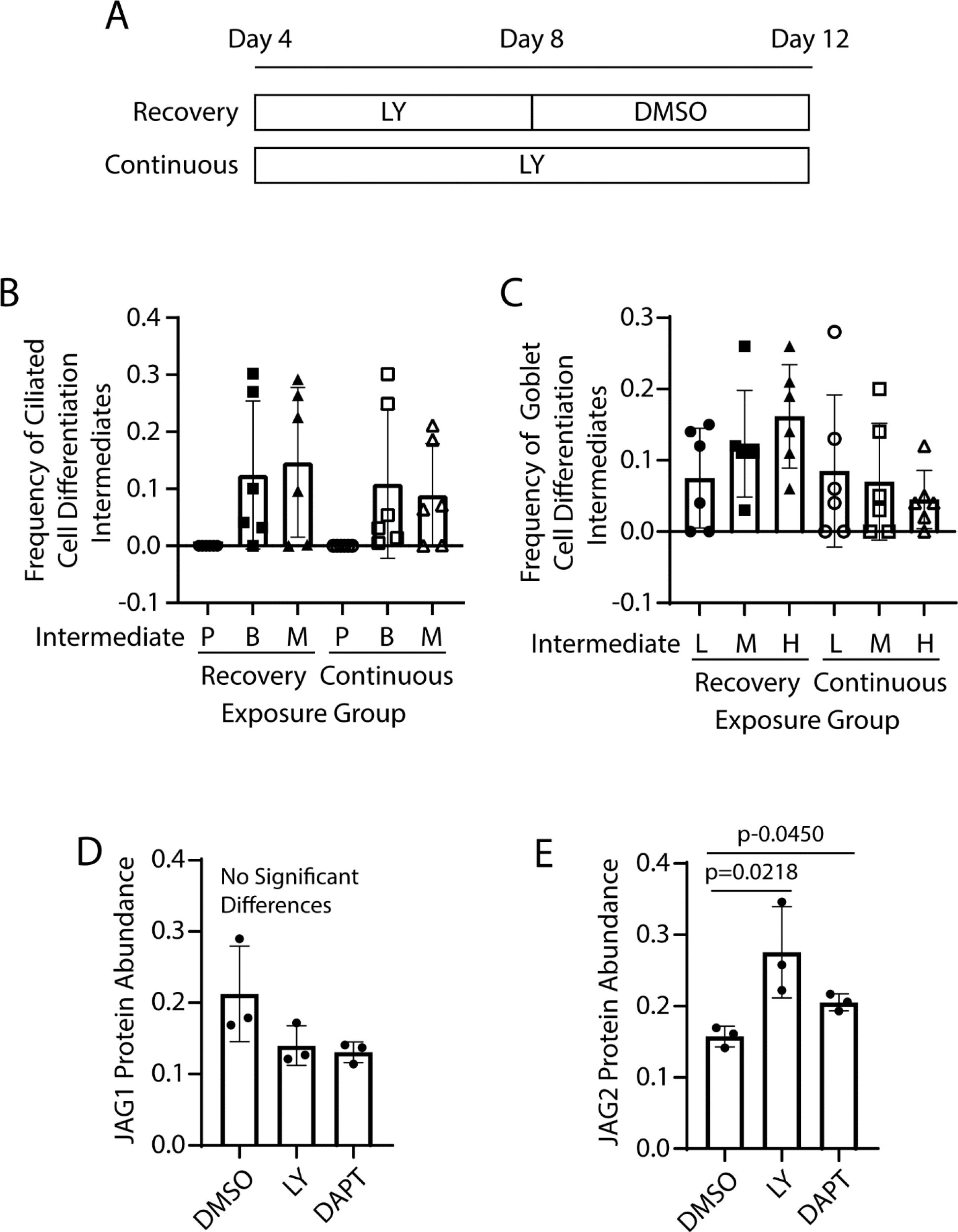
Impact of intermittent or continuous Notch inhibition of ciliated and goblet cell differentiation. **A-C**. Human bronchial basal cells were differentiated in air-liquid-interface cultures using H&H medium. Dual immunofluorescence was used to detect acetylated tubulin and MUC5B. Nuclei were stained with DAPI. (**A)** Experimental design. Recovery study: Cells were treated with 10 µM Ly on differentiation days 4 and 6 and with vehicle (DMSO) on differentiation days 8 and 10. Cultures were fixed on day 12. Continuous study: Cells were treated with 10 µM Ly on differentiation days 4, 6, 8, and 10. Cultures were fixed on day 12. (**B)** Frequency of ciliated cell differentiation intermediates. P-cells with a primary cilium, B-bristle cells, M-ciliated cells. (**C)** Frequency of goblet cell differentiation intermediates. L-MUC5B-low cells, M-MUC5B-medium cells, H-MUC5B-high cells. Mean ± SD, N=3. **D-E**. Western blot analysis of JAG1 (**D**) and JAG2 (**E**) abundance in cultures that were treated with DMSO, 10 µM LY, or 25 µM DAPT on differentiation days 8 and 10 and harvested on differentiation day 12. Mean ± SD, N=3.

### Notch regulation of ciliated and goblet cell differentiation

Previous studies demonstrated that treatment with DAPT, a γ-secretase inhibitor (GSI), inhibited Notch signaling, increased the frequency of ciliated cells, and decreased the frequency of mature goblet cells (35). To determine if this paradigm applied to the first wave of differentiation, H&H cultures were treated with DAPT on days 0-4, 2-6, 4-8, or 8-12. DAPT treatment did not alter cell density at any time point (SFig 3).

DAPT treatment significantly increased the frequency of cells with a primary cilium on day 4 (Fig 1A), significantly increased the frequency of bristle cells on days 4, 6, and 8 (Fig 1B), and significantly increased the frequency of ciliated cells on days 8 and 12 (Fig 1C). DAPT treatment did not alter the frequency of MUC5B-low or MUC5B-medium cells at any time point (Fig 1D-E). However, DAPT treatment significantly decreased the frequency of MUC5B-high cells on days 8 and 12 (Fig 1F). Similar results were gathered in experiments using a second GSI, LY411575 (LY, data not shown). Collectively, these data indicated that Notch signaling was active during the first wave of differentiation.

### Pausing of the Notch signal

Accumulation of ciliated cell differentiation intermediates suggested that GSI treatment paused the ciliation process. To investigate this mechanism, cultures were treated with LY for 4 days and recovered (recovery) or treated with LY for 8 days (continuous, Fig 2A). On differentiation day 12, ciliated and goblet cell intermediates were quantified. Cells with a primary cilium were not detected in these cultures (Fig 2B). Both bristle and ciliated cells were detected, and their frequency did not vary between the recovery and continuous treatment protocols (Fig 2B). Goblet cells defined by low, medium, and high expression of MUC5B were identified but their frequency did not vary with treatment. Since continuous Ly treatment did not cause an increase in bristle or ciliated cell frequency, these data indicated that GSI treatment paused ciliated cell differentiation.

Since Notch signaling in the tracheobronchial epithelium has been attributed to JAG1 and JAG2, the impact of GSI treatment on JAG1 or JAG2 protein abundance was determined. Cultures were treated with LY or DAPT on days 4 and 6 and protein expression was analyzed by western blot on day 8. GSI treatment did not alter JAG1 abundance (Fig 2C). In contrast, GSI treatment significantly increased the abundance of full-length JAG2 (Fig 2D). These data suggested that GSI treatment paused the differentiation process by increasing JAG2.

### Cellular localization of JAG1 and JAG2

A JAG2-dependent pausing mechanism was unanticipated and supported an in-depth analysis of JAG1 and JAG2 distribution on differentiation day 7. Immunofluorescence analysis detected JAG1 in cultures that were differentiating in each test medium (SFig 4A-C, 4J-K) and showed that JAG1 was highly localized to the perinuclear/nuclear compartments. All cells were positive for JAG1. JAG2 was also detected in all three culture conditions but was most abundant in cultures that were differentiating in H&H medium (SFig 4D-F, 4J-K). In contrast with JAG1, JAG2 was highly enriched in the cortical domain. All cells were positive for JAG2; however, expression varied from low to high.

### JAG1 subcellular location and molecular weight

The subcellular location of JAG1 on differentiation day 7 was unexpected and led to analysis at additional time points. On proliferation day 3 (Fig 3A, D, E) and differentiation day 2 (Fig 3B), JAG1 was expressed in a subset of cells and was in the perinuclear/nuclear domain. JAG1 did not colocalize with β- catenin, a component of the adherens junction (Fig 3F-G). On differentiation day 4 (Fig 3C), JAG1 was detected in most cells and some variation in protein abundance was noted.

**Figure 3:**
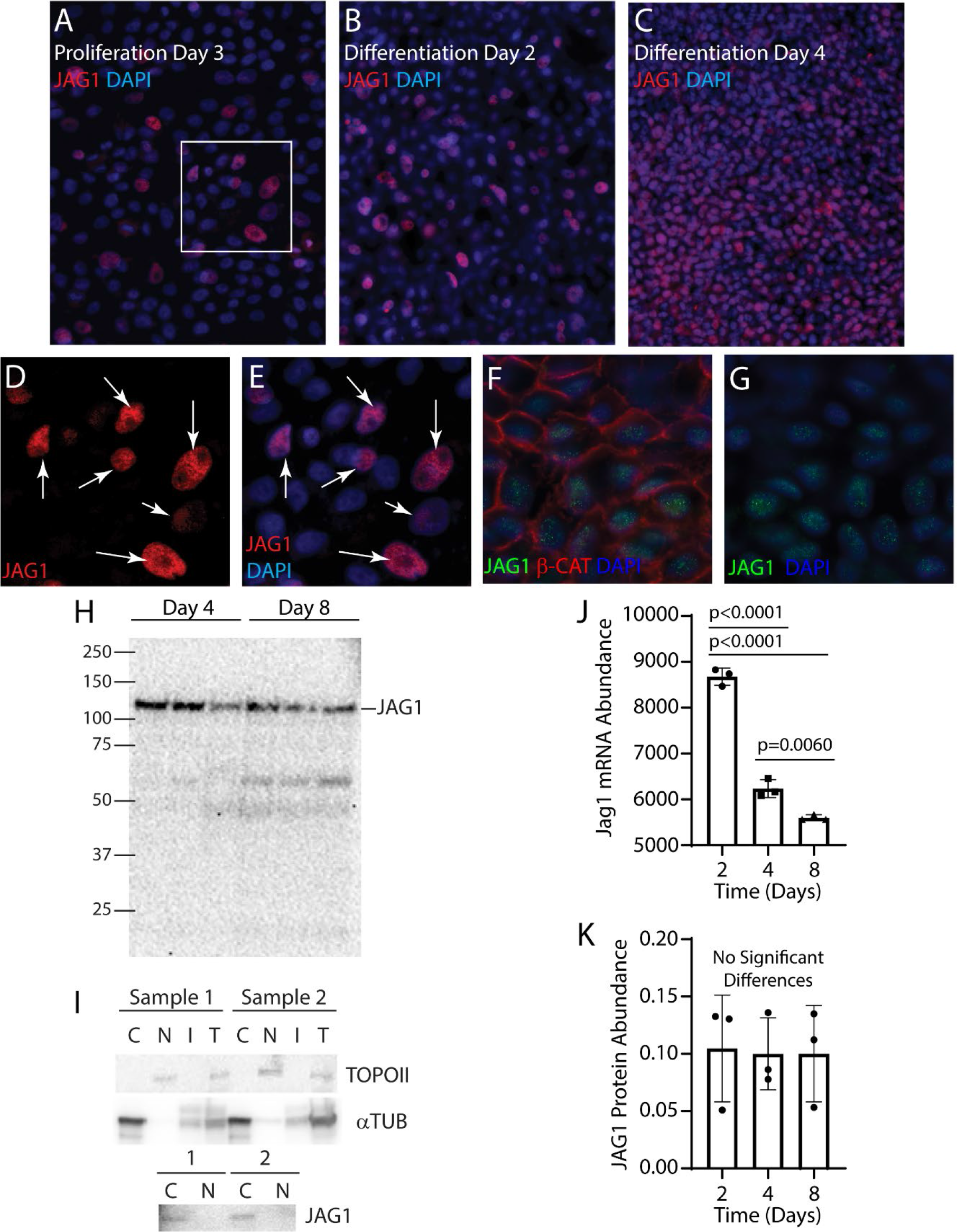
JAG1 subcellular location and length. Human bronchial basal cells were differentiated in air-liquid-interface cultures using H&H medium. **A-C.** Immunofluorescence was used to detect JAG1 (red) on proliferation day 3 (**A**), differentiation day 2 (**B**), or differentiation day 4 (**C**). Nuclei were detected with DAPI (blue). **D-E**. High magnification of the region indicated in panel A. Arrows indicate JAG1 positive cells. **F-G.** Dual immunofluorescence analysis of JAG1 (green) and β-catenin (β-Cat, red) on differentiation day 4. Nuclei were detected with DAPI (blue). Three-color (**F**) and two color (**G**) images of the same region. **H.**Western blot analysis of JAG1 on differentiation days 4 and 8. The full-length band is labeled JAG1. Three samples per time point. **I.** Cells were cultured to differentiation day 8, separated into cytoplasmic (C), nuclear (N), insoluble (I) fractions, or were unfractionated (U), and proteins were analyzed by western blot. Top panel: TOPOII, a nuclear protein; middle panel: alpha tubulin (αTUB), a cytoplasmic protein; Bottom panel: JAG1. Two different samples were analyzed. **J.** Analysis of *Jag1* mRNA abundance on differentiation days 2, 4, and 8. Mean ± SD (n=3). **K.** Western blot analysis of JAG1 protein abundance. Mean ± SD (n=3).

Previous studies reported that Notch ligands can be cleaved by γ-secretase resulting in creation of a transcriptionally active 25 kDa C-terminal peptide (51–53). Consequently, JAG1 molecular weight was determined using a JAG1 antibody which detected the C-terminal domain. On days 4 and 8, all samples contained the full-length (∼130 KDa) JAG1 protein (Fig 3H). To further examine JAG1 subcellular localization, cytoplasmic and nuclear fractions were isolated on day 8. Detection of Topoisomerase II in the nuclear fraction and alpha tubulin in the cytoplasmic fraction (Fig 3I) demonstrated successful separation. JAG1 was limited to the cytoplasmic fraction and indicated that the protein was perinuclear rather than nuclear.

Since the previous studies indicated that most if not all JAG1 was in a perinuclear compartment and that JAG1 was not acting as transcription factor, the relationship between transcription and translation was evaluated. *Jag1* mRNA was quantified by RNAseq and JAG1 protein levels were quantified by Western blot. *Jag1* abundance decreased as a function of time (Fig 3J). In contrast, JAG1 abundance was constant over time (Fig 3K). These data suggested that a post-translational mechanism regulated JAG1 abundance.

### JAG2 subcellular location and molecular weight

JAG2 localization was further evaluated on proliferation day 5 and differentiation days 4 and 8 (Fig 4A-C). The cortical pattern was observed on proliferation day 5 and JAG2-high cells were apparent on differentiation day 4. By differentiation day 8, JAG2-positive and -negative cells were noted. At each time point, JAG2 colocalized with β-catenin. On differentiation day 8, groups of cells expressing nuclear HES1 and RBPJK were detected (Fig 4D) and indicated focal Notch signaling.

**Figure 4:**
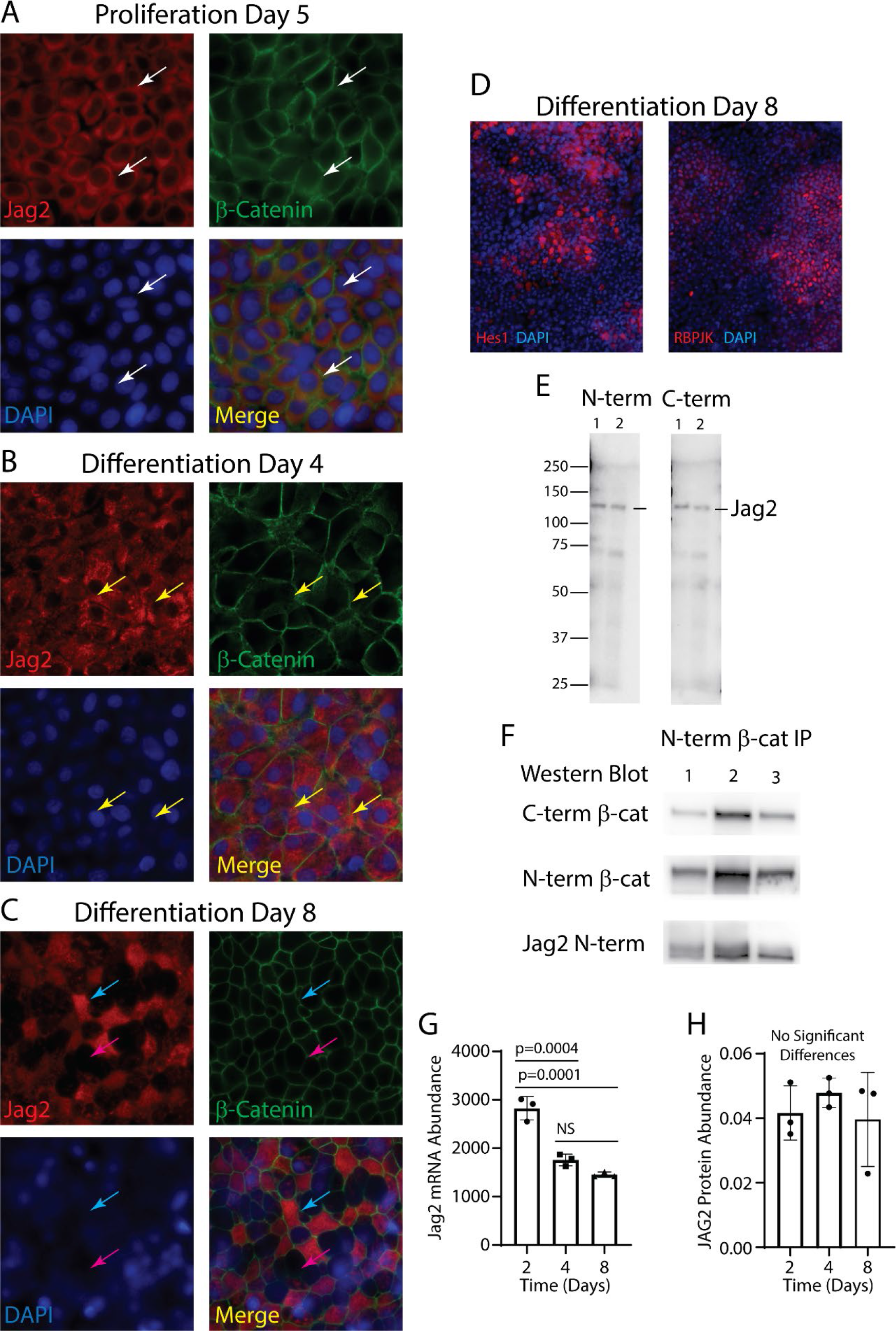
JAG2 subcellular location and length. Human bronchial basal cells were differentiated in air-liquid-interface cultures using H&H medium. **A-C.** Immunofluorescence was used to detect JAG2 (red) and β-catenin (green) on proliferation day 5 (**A**), differentiation day 4 (**B**), or differentiation day 8 (**C**). Nuclei were detected with DAPI (blue). Each panel presents single color and merged images. Arrows: white, cortical JAG2; yellow, JAG2-high cells; blue, JAG2- positive cells; pink, JAG2-negative cells. **D.** Immunofluorescence analysis of HES1 (red) and RBPJK (red) on differentiation day 8. Nuclei were detected with DAPI (blue). **E.** Western blot analysis of JAG2 on differentiation days 4 (lane 1) and 8 (lane 2). The full-length band is labeled JAG2. **F**. Co-immunoprecipitation analysis of JAG2 and β-catenin on differentiation day 8. Proteins were immunoprecipitated with an N-terminus (N-term) specific β-catenin antibody. Precipitates were analyzed for β-catenin using a C-terminus (C-term) specific β-catenin antibody, N-term specific β-catenin antibody, or a JAG2 N-term specific antibody. Three samples were analyzed. **G.** Analysis of *Jag2* mRNA abundance on differentiation days 2, 4, and 8. Mean ± SD (n=3). **H.** Western blot analysis of JAG2 protein abundance. Mean ± SD, N=3. **G.**

The molecular weight of JAG2 was determined using two antibodies which were specific to the N-terminal or C-terminal domains. This study detected full-length (∼145 KDa) JAG2 on days 4 and 8 (Fig 4E). To further examine JAG2 subcellular localization, co-immunoprecipitation was used to determine if JAG2 was associated with the adherens junction. An N-terminal specific β- catenin antibody was used for immunoprecipitation and the precipitate was analyzed with N-terminal and C-terminal specific β-catenin antibodies. β-catenin was successfully isolated (Fig 4F) and JAG2 was identified in the β-catenin complex.

Since the previous studies suggested that much of JAG2 was trafficking to or from the plasma membrane, the relationship between transcription and translation was evaluated using RNAseq and western blots. *Jag2* abundance decreased as a function of time (Fig 4G). In contrast, JAG2 abundance was constant over time (Fig 4H). Like JAG1, these data suggested a post-translational mechanism regulated JAG2 abundance.

### Regulation of JAG2 abundance

Identification of JAG2-positive and JAG2-negative cells on differentiation day 8 suggested that JAG2 was degraded in some cells. A candidate regulator is the WNT/β-catenin pathway (35). To examine this mechanism, ALI cultures were treated with various concentrations of CHIR99021 (CHIR, a WNT/β-catenin agonist, (54)) or XAV939 (XAV, a WNT/β-catenin antagonist, (55)) on differentiation days 4 and 6 and cell lysates were collected on day 8. Western blot analysis demonstrated that neither CHIR nor XAV treatment altered the abundance of JAG1 (Fig 5A). In contrast, both treatments caused a significant decrease in JAG2 abundance (Fig 5B). Interestingly, the CHIR effect was limited to lower doses (5 and 10 µM); whereas the XAV effect was observed at all three doses.

**Figure 5:**
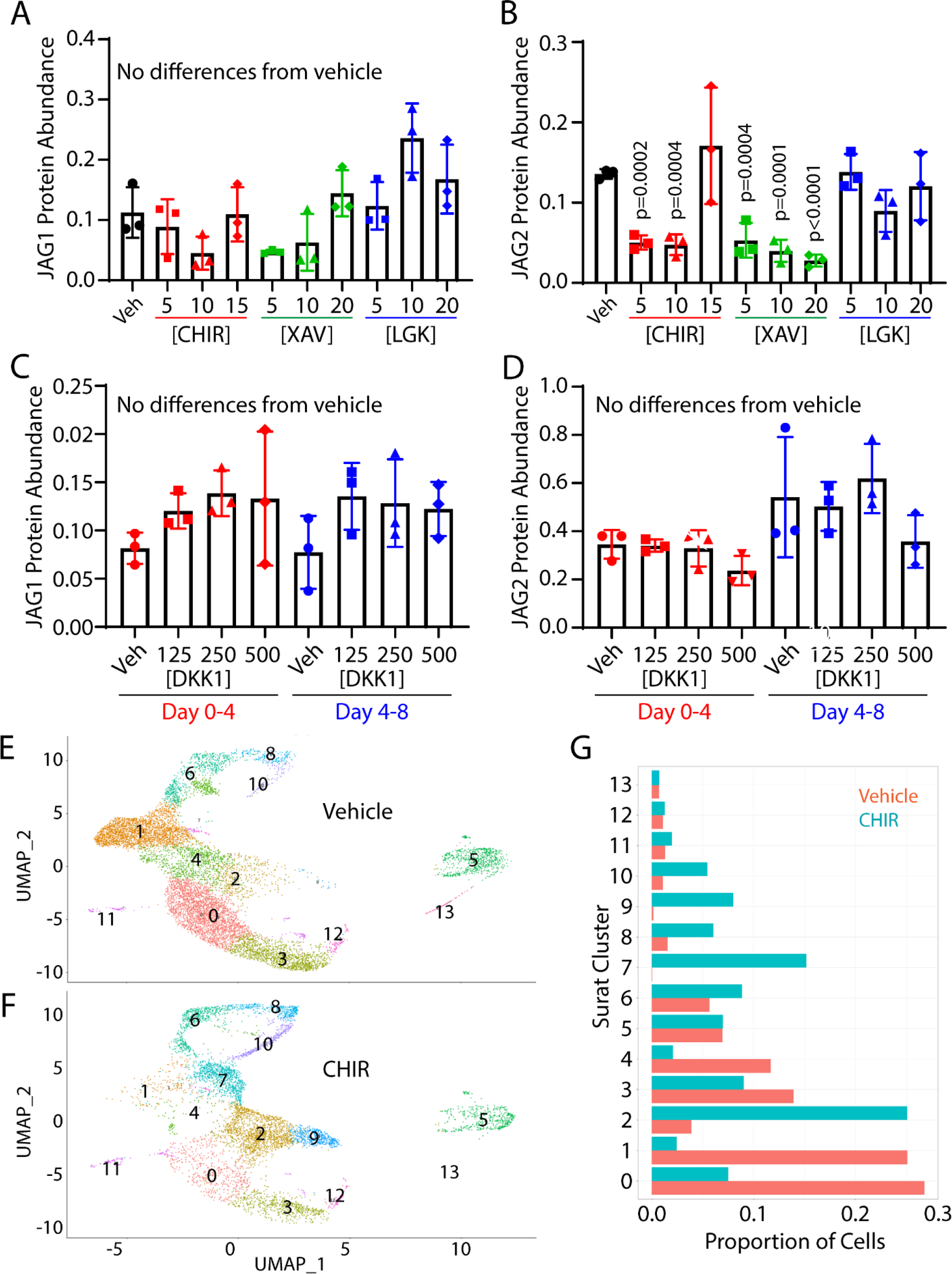
Regulation of JAG1 and JAG2 abundance. Human bronchial basal cells were differentiated in air-liquid-interface cultures using H&H medium. **A-B.** On differentiation days 4 and 6, the cultures were treated with vehicle, (Veh, DMSO), CHIR99021 (CHIR), XAV939 (XAV), or LGK974 (LGK) and protein lysates were collected on day 8. Drug concentrations are in µM. Western blots were used to quantify JAG1 (**A**) and JAG2 (**B**) abundance. Mean ± SD, N=3. **C-D.** Cultures were treated with Veh or DKK1 on differentiation days 0 and 2 and lysed on day 4 or treated on differentiation days 4 and 6 and lysed on day 8. Drug concentrations are in ng/ml. Western blots were used to quantify JAG1 (**C**) and JAG2 (**D**) abundance. Mean ± SD, N=3. **E-F**. Uniform manifold approximation and projection (UMAP) analysis of cells that were treated with vehicle (**E**) or CHIR (**F**) on days 4 and 6 and harvested for scRNAseq analysis on day 8. **G.** Number of normalized cells per Surat Cluster.

### Impact of CHIR and XAV on the transcriptome

CHIR and XAV are typically thought to regulate WNT/β-catenin pathway by altering gene expression. To evaluate this mechanism, ALI cultures were treated with vehicle, 10 µM CHIR or 10 µM XAV on differentiation days 4 and 6 and RNA was purified on day 8. RNA-sequencing and Gene Set Expression Analysis (GSEA) demonstrated that CHIR upregulated expression of WNT/β-catenin target genes including *Axin2*, *Lef1*, *Tcf7* (SFig 5A-B, SFig 6). These genes contributed to the GSEA Leading-Edge and had a high impact on the GSEA Enrichment Score (an indicator of changes in pathway activity). Other GSEA Leading-Edge genes were *Wnt5B* and known pathway antagonists (*Axin1, Dkk1*, *Dkk4,* and *Nkd1*). Since these genes and *Axin2* inhibit the WNT/β-catenin pathway and *Lef1* and *Tcf7* repress gene expression, these data suggested that CHIR antagonized the WNT/β- catenin pathway on day 8.

In keeping with its perceived function as a WNT/β-catenin pathway antagonist, XAV treatment downregulated expression of *Axin2*, *Lef1*, and *Tcf7*, and *Wnt5B* (SFig 5C-D, SFig 6). However, most GSEA Leading-Edge WNT/β-catenin pathway genes were typical of the Notch pathway (*Dll1*, *Hey1*, *Jag1*, *Jag2*, *Notch1*, and *Notch4*). Analysis of the Notch gene set indicated that XAV treatment caused a significant downregulation of Notch signaling (SFig 5G-H). A similar analysis of the Notch gene set in CHIR treated cultures also indicated downregulation (SFig 5E- F). Although the high FDR q-value suggested cautious interpretation of the CHIR-Notch pathway interaction, many of the GSEA Leading-Edge genes were shared between the CHIR and XAV groups (*Dll1*, *Jag1*, *Notch1*, *Lfng*, *Prkca*, *Psenen*, *Sap30*, and St3gal6, SFig 5E-H, SFig 6).

Although *Jag1* and *Jag2* contributed to the Enrichment Scores, neither gene product was consistently included in the GSEA Leading-Edge gene sets. Consequently, the impact of CHIR and XAV treatment on gene expression was evaluated. *Jag1* and *Jag2* abundance did not vary by treatment (SFig 5I-J). Collectively, the gene expression data indicated that the main effect of CHIR or XAV treatment was to antagonize WNT/β-catenin and Notch signaling and that decreased signaling could not be attributed to a change in *Jag1* or *Jag2* gene expression.

### WNT independent regulation of JAG2 abundance

To examine the impact of WNT antagonism on JAG1 and JAG2 abundance, ALI cultures were treated with vehicle, LGK974 (LGK) a porcupine inhibitor which prevents WNT secretion, or DKK1 which inhibits WNT signaling by binding the LRP5/6 coreceptor. Neither LGK treatment (Fig 5A-B) nor DKK1 treatment (Fig 5C-D) altered JAG1 or JAG2 abundance. These data suggested that a WNT-independent mechanism regulated the abundance of JAG2 in CHIR and XAV treated cultures.

### CHIR regulation of cell fate

While the decrease in JAG2 abundance in response to CHIR or XAV treatment was unexpected, a previous study reported that both drugs altered secretory and ciliated cell differentiation (35). To further examine changes in cell fate, ALI cultures were treated with CHIR on days 4 and 6 and cells were isolated for single cell RNAseq in day 8.

Fourteen clusters (0-13) were identified (Fig 5E, F) and 10 clusters differed significantly between vehicle and CHIR treated cultures (Fig 5G). In vehicle treated controls, clusters 6, 8, and 10 exhibited a mitotic signature (STable 1). The frequency of cells in these clusters increased in CHIR treated cultures. Cluster 1 was prominent in vehicle treated cultures and contained basal cells which exhibited a matrix production signature. This cluster over-expressed *Jag2* and *Wnt4* and was nearly absent in CHIR treated cultures. Clusters 4, 0, 3 and 12 contained cells within the secretory lineage and clusters 4, 0 and 3 expressed Hes4, Hes1, and/or Hes2. Cluster 3 contained secretory primed basal cells that were defined by expression of *Ceacam6*, *Scgb1A1*, and *Scgb3A2*. Cells within cluster 3 were less mature than those in cluster 12. Clusters 4, 0 and 3 were significantly decreased in CHIR treated cultures. Clusters 13 and 5 contained ciliated cell lineage cells. Cluster 13 was enriched in early-stage ciliated cells defined by expression of *Mcidas* and cluster 5 contained more mature ciliated cells. Cluster 2 was increased in CHIR treated cultures and clusters 7 and 9 were unique to CHIR treated cultures. Cluster 7 was distinguished by down-regulation of WNT pathway antagonists (*Axin2*, *Dkk2*, *Dkk4*, *Nkd1*), and the presence of cell motility and keratinization signatures. The keratinization signature was upregulated in clusters 2 and 9. Since keratinization was previously associated with squamous differentiation in the tracheobronchial epithelium (56), these data indicated that CHIR treatment promoted this fate and suggested that JAG2 suppressed squamous differentiation.

### JAG1 and JAG2 trafficking to the cell surface

To further investigate roles for JAG1 and JAG2 in Notch signaling, cell surface protein was evaluated by immunofluorescence analysis of non-permeabilized cultures. JAG1 was not detected on the cell surface on differentiation day 8. In contrast, JAG2 was detected on the surface of many cells (Fig 6A-B). Subsequent permeabilization and staining detected both JAG1 and JAG2 and illustrated their typical intracellular distributions (Fig 6C-F).

**Figure 6:**
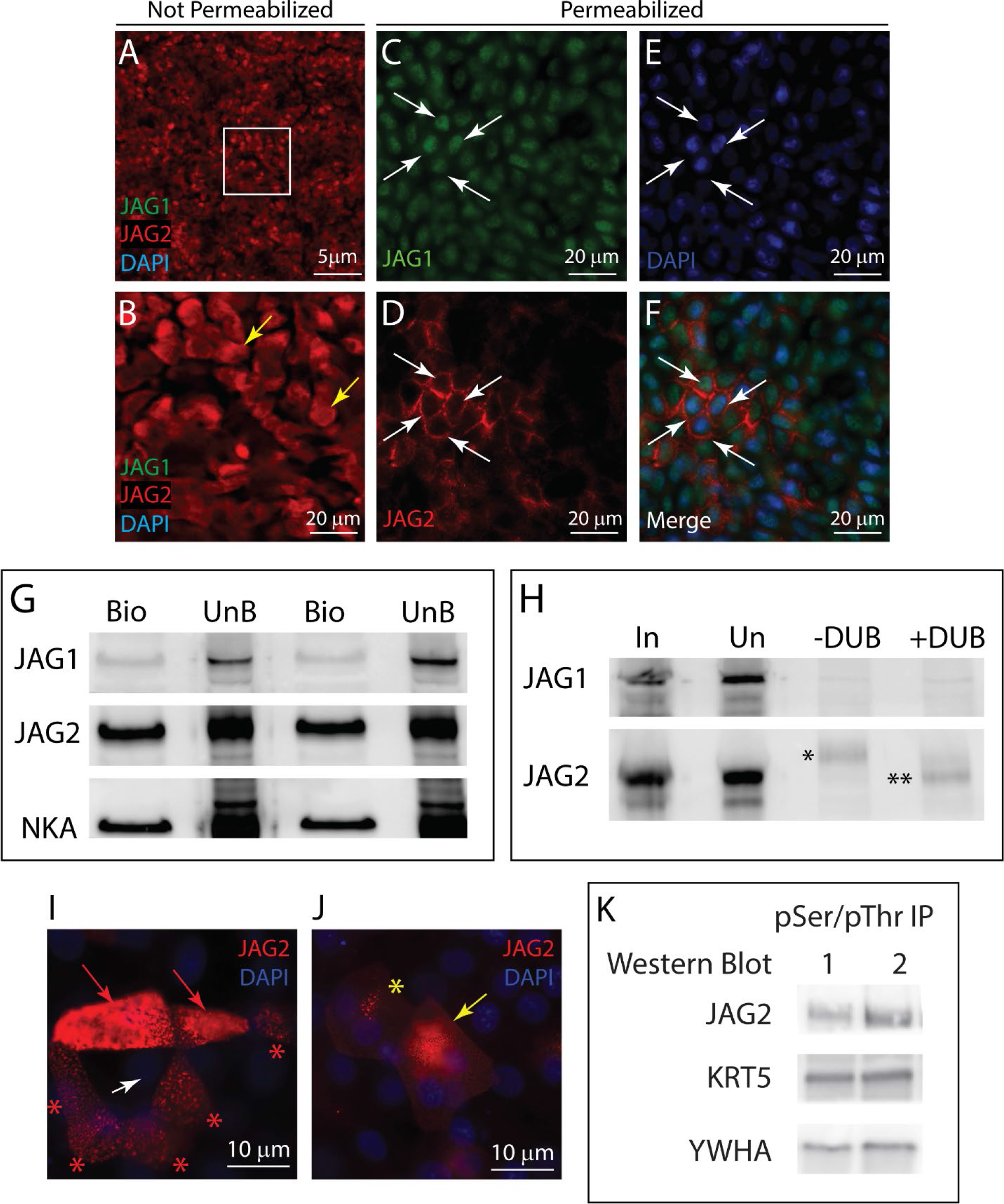
JAG1 and JAG2 trafficking to the cell surface. Human bronchial basal cells were differentiated in air-liquid-interface cultures using H&H medium. **A-B**. On differentiation day 8, cultures were fixed under non-permeabilization conditions and JAG1 (green) and JAG2 (red) were detected by dual immunofluorescence staining. Nuclei were stained with DAPI (blue). Panel B is a higher magnification of the region indicated in panel A. **C-F.** After imaging, the cultures were permeabilized and re-stained for JAG1 (green), JAG2 (red), and DAPI (blue). Arrows: antigen-positive cells. **G.** On differentiation day 8, cell surface proteins were labeled with biotin, biotinylated proteins were recovered, and the bound (BIO) and unbound (UnB) fractions were evaluated by western blot. Two samples were evaluated. **H.**On differentiation day 8, cell lysates were reacted with TUBE beads which bind to proteins decorated with 4 or more ubiquitin moieties. The bound fraction was treated with vehicle (-DUB) or a broad spectrum debubiquitinase (+DUB). Western blots were generated from the unfractionated (In), unbound (Un), -DUB, and +DUB samples and were evaluated for JAG1 and JAG2. *Ubiquitinated JAG2, **deubiquitinated JAG2. Immunofluorescence was used to detect JAG2 in Veh (**I**) and CHIR (5 µM, **J**) treated cultures on differentiation day 8. Arrows: red, JAG2-high cell; white, JAG2-negative cell; yellow, redistributed JAG2. Asterisks: JAG2-low cells. **K.** Cells were cultured to differentiation day 8, lysed, and immunoprecipitated with a phospho-serine/phospho- threonine (pSer/pThr) antibody. Precipitates were analyzed for JAG2 using the C-terminus specific antibody, KRT5 antibody, or a pan-YWHA antibody. Two samples were analyzed.

Cell surface biotinylation was used to evaluate JAG1 and JAG2 trafficking to the plasma membrane. These studies detected sodium/potassium ATPase (NKA) a known cell surface protein as well as JAG1 and JAG2 (Fig 6G). However, cell surface JAG2 was 5-10 times more abundant than JAG1. Since interaction with a Notch receptor results in ligand ubiquitination and internalization, polyubiquitination of JAG1 and JAG2 was analyzed. Little or no JAG1 was detected in the bound (ubiquitin-positive) fraction (Fig 6H, upper panel). In contrast, JAG2 was detected in the bound fraction and the molecular weight of the captured JAG2 decreased after treatment with a deubiquitinase (Fig 6H). These biochemical studies indicated that JAG2 participated in Notch-signaling and raised the possibility that JAG1 was not involved.

### Phosphorylation of JAG2

Immunofluorescence analysis demonstrated that CHIR treatment caused a striking redistribution of JAG2 to a perinuclear location (Fig 6I-J) and was suggestive of JAG2 degradation. JAG2 was not detected by immunofluorescence staining in XAV treated cells (data not shown). CHIR is a potent and highly selective inhibitor of GSK3 (54). In contrast, XAV inhibits tankyrase and regulates GSK3 by altering the kinase’s localization within the cell (55). Since both drugs decrease GSK3 activity and decrease JAG2 abundance, it was possible that the shared mechanism was a GSK3-dependent decrease in JAG2 phosphorylation. To evaluate this mechanism, differentiation day 8 cell lysates were immunoprecipitated with an anti-pSerine/pThreonine (pS/pT) antibody and western blots were probed for JAG2 and positive controls (KRT5 and YWHA). This study detected JAG2, KRT5 and YWHA in the bound fraction (Fig 6K) and indicated that JAG2 was phosphorylated on Serine and/or Threonine. Collectively, these data suggested that normal JAG2 trafficking was dependent on phosphorylation by GSK3.

### Roles for JAG1 and JAG2 in Notch Signaling

To determine if JAG1 was involved in Notch signaling, cultures were treated with a JAG1 DSL-domain peptide on differentiation days 8 and 10 and cultures were fixed on differentiation day 12. Immunofluorescence analysis of ACT and MUC5B was used to detect differentiation intermediates. Cells expressing a primary cilium were not detected (Fig 7A). However, vehicle treated cultures contained a small number of bristle cells and many ciliated cells. Treatment with the JAG1 peptide caused a significant increase in the frequency of bristle cells but did not alter the frequency of ciliated cells (Fig 7A). The same cultures did not contain MUC5B-low cells (Fig 7B). However, MUC5B-medium, and MUC5B- high cells were detected. Treatment with JAG1 peptide did not alter the frequency of MUC5B- medium cells but significantly decreased the frequency of MUC5B-high cells. These data suggested two JAG1-dependent processes. First, that the JAG1 peptide inhibited Notch signaling resulting in increased basal-to-bristle cell differentiation. Second, the decreased maturation of MUC5B-medium cells combined with the increase in bristle cell frequency raised the possibility that JAG1 peptide treatment promoted MUC5B-medium to ciliated cell differentiation.

**Figure 7:**
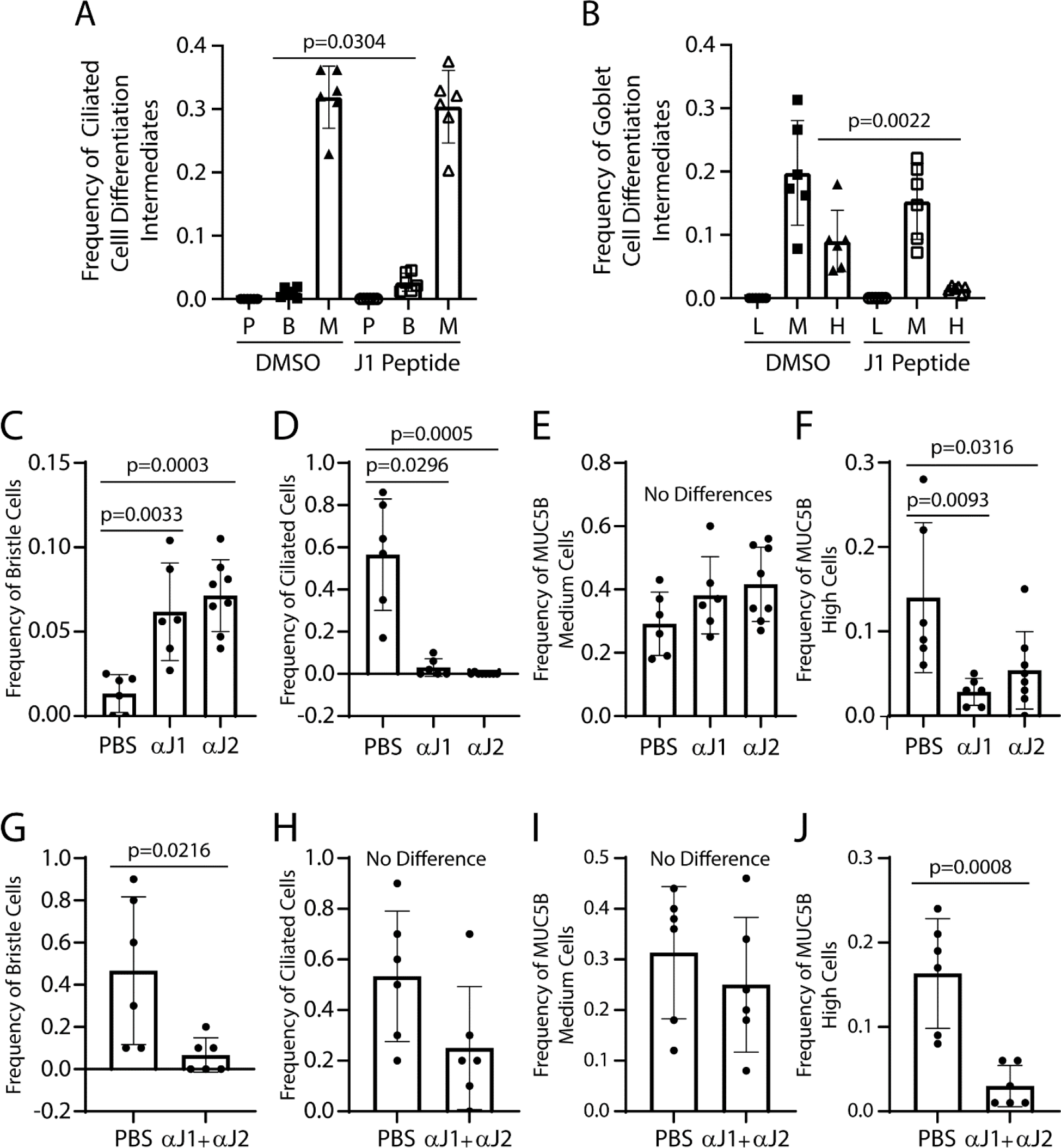
Roles for JAG1 and JAG2 in ciliated and goblet cell differentiation. Human bronchial basal cells were differentiated in air-liquid-interface cultures using H&H medium. Treatments were on differentiation days 8 and 10 and cultures were fixed on day 12. Differentiation was quantified by immunofluorescence analysis of acetylated tubulin, MUC5B, and DAPI. **A-B.** Cells were treated with vehicle (DMSO) or 40 µM JAG1 peptide. Frequency of ciliated cell differentiation intermediates. P-cells with a primary cilium, B-bristle cells, M-ciliated cells (**A**). Frequency of goblet cell differentiation intermediates (**B**). L-MUC5B-low cells, M-MUC5B- medium cells, H-MUC5B-high cells. **C-F.** Cells were treated with vehicle (PBS) or 25 µg/ml neutralizing antibody to JAG1 (αJ1) or JAG2 (αJ2). Frequency of bristle cells (**C**). Frequency of ciliated cells (**D**). Frequency of MUC5B-medium cells (**E**). Frequency of MUC5B-high cells (**F**). **G-J.** Cells were treated with PBS or 25 µg/ml αJ1 and 25 µg/ml αJ2. Frequency of bristle cells (**G**). Frequency of ciliated cells (**H**). Frequency of MUC5B-medium cells (**I**). Frequency of MUC5B-high cells (**J**). Mean ± SD, N=6.

Since the JAG1 peptide is 71% identical to the JAG2-DSL domain, it was unclear if the JAG1 peptide results could be attributed to JAG1. Further, the JAG1 peptide results were consistent with reports that used JAG1 peptide to antagonize Notch signaling (57) but disagreed with those reporting that this reagent agonized Notch signaling (58, 59). Consequently, additional studies used neutralizing antibodies to inhibit JAG1 or JAG2 activation of Notch receptors (29).

To address the limitations of the JAG1 peptide studies, ALI cultures were treated with neutralizing antibodies on days 8 and 10 and ciliated and secretory cell differentiation was assayed on day 12. Treatment with anti-JAG1 or anti-JAG2 significantly increased bristle cell frequency and decreased ciliated cell frequency (Fig 7C-D). Neutralizing antibody treatment did not alter the frequency of MUC5B-medium cells but significantly decreased the frequency MUC5B-high cells (Fig 7E-F). These data suggested that loss of either JAG1 or JAG2 caused MUC5B-medium cells to generate bristle cells. To address the finding that anti-JAG1 or anti- JAG2 treatment decreased ciliated cell frequency, ALI cultures were treated with a mixture of anti-JAG1 and anti-JAG2. This treatment significantly decreased bristle cell frequency but did not alter ciliated cell frequency (Fig 7G-H). The combined neutralizing antibody treatment did not alter the frequency of MUC5B medium cells and significantly decreased the frequency of MUC5B-high cells (Fig 7I-J). These data in combination with the single antibody study indicated that either JAG1 or JAG2 inhibited bristle cell to ciliated cell differentiation.

## DISCUSSION

This study demonstrated that JAG1 and JAG2 cooperate to determine the fate of the MUC5B- medium (secretory progenitor) cell (SFig 7A). The neutralizing antibody studies found that both JAG1 and JAG2 were required to generate a mature MUC5B-high goblet cell. In contrast, neutralization of either JAG1 or JAG2 allowed the MUC5B-medium cell to generate a bristle cell. Finally, the CHIR and XAV studies indicated that degradation of JAG2 resulted in a squamous fate (SFig 7B). While these data support the paradigm that high Notch signal strength selected the goblet cell fate, the possibility of moderate and low Notch intensity within the secretory lineage was novel.

Previous studies indicated that cell fate was determined by different assemblies of Notch receptors (13, 20). The present study suggests that different assemblies of JAG ligands determine Notch signal intensity. First, JAG1 was detected in all differentiating cells. Second, a striking change in cell morphology coincided with the emergence of JAG2 positive and negative cells and raised the possibility that increased polarization concentrated JAG2 at cell-cell junctions (60). When combined with the histological and biochemical data, these studies indicated that JAG1 might establish a minimum ligand concentration while changes JAG2 trafficking result in cells that express high or low levels of cell surface ligand. Thus, it is possible that the MUC5B-medium cell fate is determined by its location in a ligand-low (JAG1 only), ligand-medium (mixture of JAG1+/JAG2+ and JAG1+/JAG2- cells), or a ligand-high (JAG1+/JAG2+) environment (Graphical Abstract).

This study also indicated that JAG2 paused the ciliated cell differentiation process (SFig 7C). Initial support for pausing came from the finding that GSI treatment increased the frequency of ciliated cell differentiation intermediates. These data suggested that Notch signaling fluctuated from “off” to “on” as the cell progressed from one morphological intermediate to the next. These data challenged the idea that ciliated differentiation occurred in the absence of a Notch signal but also raised the possibility that the GSI-sensitive target was the ligand rather than the receptor. Support for the latter idea comes from previous demonstration that JAG1 and JAG2 were cleaved by γ-secretase resulting in production of an intracellular domain which functions as a transcription factor (51–53). Failure to detect the ∼25 kDa intracellular domain may reflect low sensitivity in the ALI model relative to ligand over-expression systems.

Evidence of pausing also challenged the notion that ciliated cell differentiation was continuous and suggested that differentiation occurred in incremental steps. Demonstration that expression of a ciliated transcription network gene product, *cMyb,* fluctuated over time and regulated progression through the ciliated cell differentiation program (61) supported this interpretation.

Although the neutralizing antibody studies indicated that bristle-to-ciliated cell differentiation occurred only when JAG1 and JAG2 were blocked, western blot analysis correlated the GSI- dependent signal with an increase in full-length JAG2 abundance. Since gene expression studies indicated that GSI treatment did not alter *Jag2* (or *Jag1*) mRNA abundance, it is possible that pausing involved a change in JAG2 protein trafficking.

Previous studies showed that transmembrane proteins, including JAG1 and JAG2, transit the endocytic recycling compartment as they move to the plasma membrane. This compartment is positioned under the nucleus in adherent cells, and sorts endocytosed proteins that are recycled to the plasma membrane, enter the lysosome for degradation, or are sequestered in a perinuclear site termed the pericentrion (62). A protein’s trafficking pattern is determined in part by phosphorylation and ubiquitination and changes in the rate of movement through the various endosomal compartments regulate Notch signaling (63).

This study identified a striking difference in JAG1 and JAG2 localization within the cytoplasm of differentiating cells. Cell fractionation studies indicated that JAG1 was highly enriched in the perinuclear domain. Demonstration that JAG1 was not poly-ubiquitinated indicated that most JAG1 was in the endocytic recycling compartment and not sequestered in the pericentrion. In contrast, the subcellular distribution and polyubiquitination of JAG2 were consistent with endocytosis, recycling through the endocytic pathway, and post-signaling degradation via the lysosome. Increased JAG2 protein levels in GSI-treated cultures and decreased JAG2 levels in CHIR and XAV treated cultures suggested that JAG2 routing could be altered and that JAG2 and GSK3 might be co-localized in multivesicular bodies (64, 65).

A previous study reported that NOTCH1 trafficking to the cell surface was inhibited by GSK3- mediated phosphorylation (66) while the present study showed that GSK3 inhibition resulted in JAG2 degradation. Consequently, GSK3 activity might determine if a cell traffics JAG2 or NOTCH1 to the cell surface. Although the WNT/β-catenin pathway was a likely regulator of GSK3 activity, our previous report found that CHIR and XAV (35) inhibited ciliated cell and goblet cell commitment and suggested a complex mechanism. The present study indicates that both CHIR and XAV antagonized WNT/β-catenin signaling and that the main effect of these drugs was downregulation of Notch signaling. Although these data suggest that Notch signaling is dependent on β-catenin, a role for WNT signaling was discounted by studies which inhibited WNT secretion or WNT interaction with LRP6/5. A similar WNT ligand-independent role for GSK3 in cell fate specification was previously reported for iPSC that were undergoing directed differentiation (67) and suggested that establishment of tracheobronchial ligand- and receptor-expressing cells was regulated by a GSK3-dependent and WNT-independent mechanism.

## CONCLUSION

This study demonstrates that JAG1 and JAG2 are necessary for cell fate specification within the tracheobronchial epithelium. Differences in JAG1 and JAG2 trafficking raise the possibility that distinct assemblies of these ligands generate high, medium, and low Notch signaling environments within the differentiating epithelium (Graphical Abstract). Further analysis of JAG trafficking has the potential to identify treatments which could normalize goblet and ciliated cell frequency in people who have chronic mucosecretory lung diseases and those whose ciliated cell population is depleted by exposure to environmental agents.

## ACKNOWLEDGEMENTS

This project was funded by a Research Grant from Cystic Fibrosis Foundation (REYNOL17XX0- 20XX0, SDR), the Nationwide Children’s Hospital Cell Based Therapy Program (47306-0002- 1217, SDR), and a Nationwide Children’s Hospital Technology Development Award (Technology 2019-015, SDR). We acknowledge the Abigail Wexner Research Institute Center for Genomics Research which conducted the RNA sequencing and analysis, and the C3 RDP which provided human bronchial cells (MCCOY19R0). JAG1 and JAG2 neutralizing antibodies were provided by Genentech, Inc. under a Materials Transfer Agreement.

## Author Contributions

SDR: Conception and design, acquisition of laboratory data, interpretation of data, manuscript preparation and review.

CLH: Acquisition of laboratory data, manuscript review

AA: Acquisition of laboratory data, manuscript review

SWL: Acquisition of laboratory data, manuscript review

SW: RNAseq and scRNAseq analysis, manuscript review

ZHT: scRNAseq analysis, manuscript review

TC: Data review and interpretation, manuscript review

ECB: Data review and interpretation, manuscript review

---

**Supplemental Figure 1:**
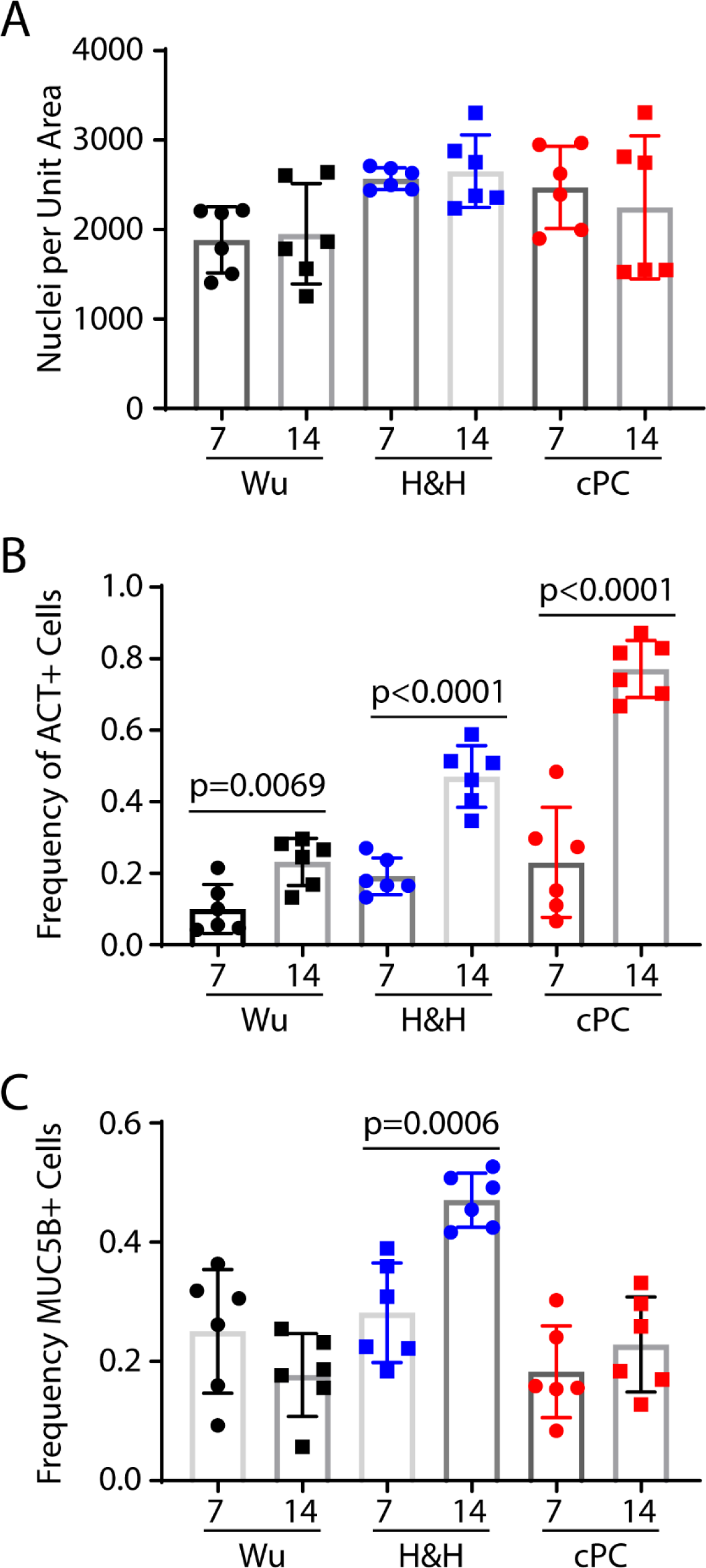
Ciliated and goblet cell differentiation as a function of medium type. Human bronchial basal cells were differentiated in air-liquid-interface cultures using three media: Wu, H&H (H&H), and complete Pneumacult (cPC). Cultures were fixed on days 7 and 14. **A.** Cell density was determined by quantifying the number of DAPI-stained nuclei per unit area. **B.** Ciliated cells were identified by acetylated tubulin (ACT) staining and their frequency was reported as the number ACT+ cells/number nuclei. **C.** Goblet cells were identified by MUC5B staining and their frequency was reported as the number MUC5B+ cells/number nuclei. Mean ± SD, N=6.

**Supplemental Figure 2:**
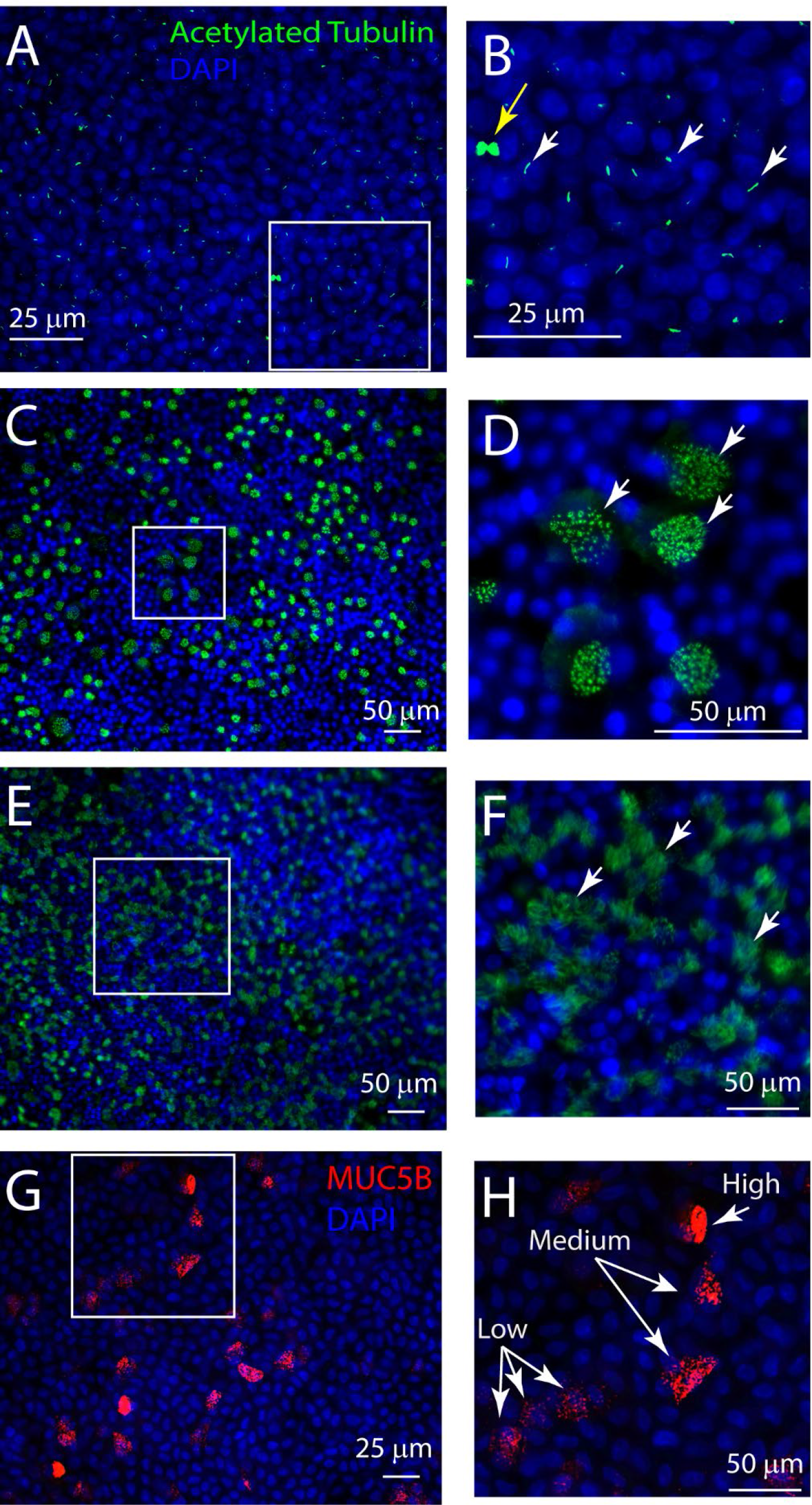
Intermediate ciliated and goblet cell phenotypes. Human bronchial basal cells were differentiated in air-liquid-interface cultures using H&H medium. **A-B.** Cells defined by a primary cilium were identified by ACT staining on differentiation day 4. Panel B is a high magnification image of the region identified in panel A. Arrows: white, primary cilium; yellow, mitotic figure. **C-D.** Cells with bristle morphology were identified by ACT staining on differentiation day 8. Panel D is a high magnification image of the region identified in panel C. Arrows: bristle cells. **E-F.** Ciliated cells defined were identified by ACT staining on differentiation day 12. Panel F is a high magnification image of the region identified in panel E. Arrows: Ciliated cells. G-H. Goblet cells were identified by MUC5B staining on differentiation day 8. Panel H is a high magnification image of the region identified in panel G. Arrows identify MUC5B-low, -medium, and -high goblet cell intermediates.

**Supplemental Figure 3:**
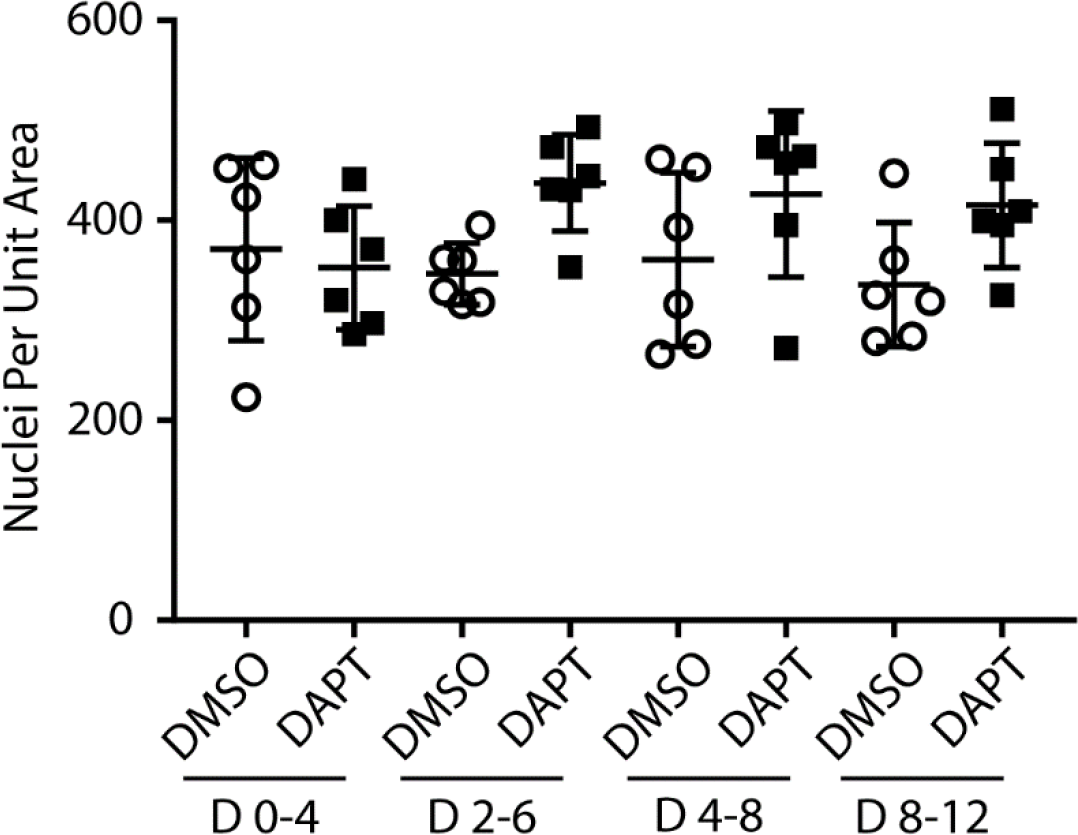
Cell density in vehicle and DAPT treated cultures. Human bronchial basal cells were differentiated in air-liquid-interface cultures using H&H medium. Cells were treated with vehicle (DMSO) or 25 µM DAPT as follows: treatment on differentiation days 0 and 2 and fix on day 4, treatment on differentiation days 2 and 4 and fix on day 6, treatment on differentiation days 4 and 6 and fix on day 8, or treatment on differentiation days 8 and 10 and fix on day 12. Nuclei were stained with DAPI and quantified. Mean ± SD, N=6.

**Supplemental Figure 4:**
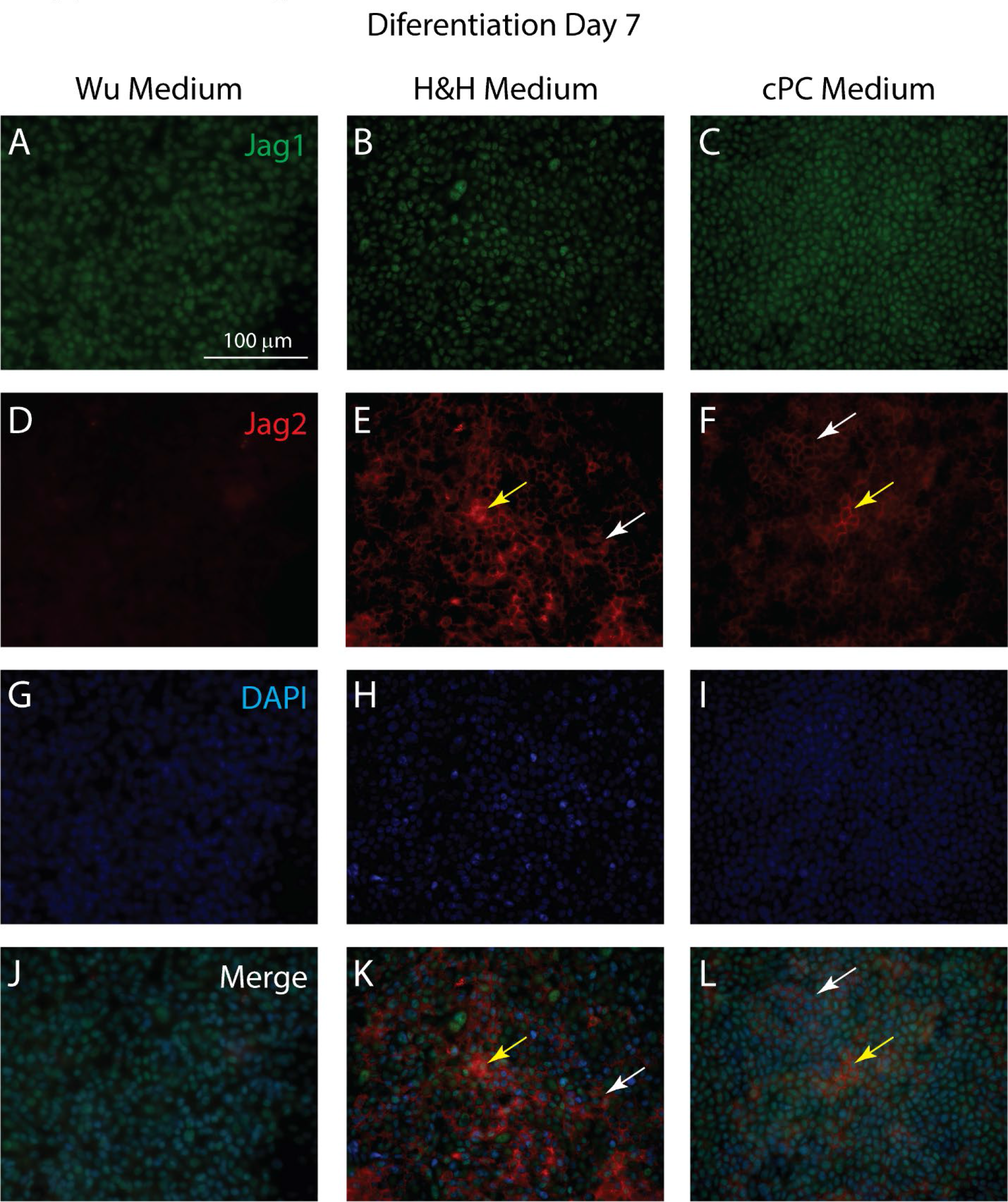
JAG1 and JAG2 localization as a function of medium type. Human bronchial basal cells were differentiated in air-liquid-interface cultures using three media: Wu (**A, D, G, J**), H&H (H&H, **B, E, H, K**), and complete Pneumacult (cPC, **C, F, I, L**). Cultures were fixed on day 7. JAG1 (green, **A-C**) and JAG2 (red, **D-F**) were detected by dual immunofluorescence. Nuclei were detected with DAPI (blue, **G-I**). Merged images (**J-L**). Arrows: yellow, JAG2 expression; white, low JAG2 expression. Scale bar in panel A, 100 µm.

**Supplemental Figure 5:**
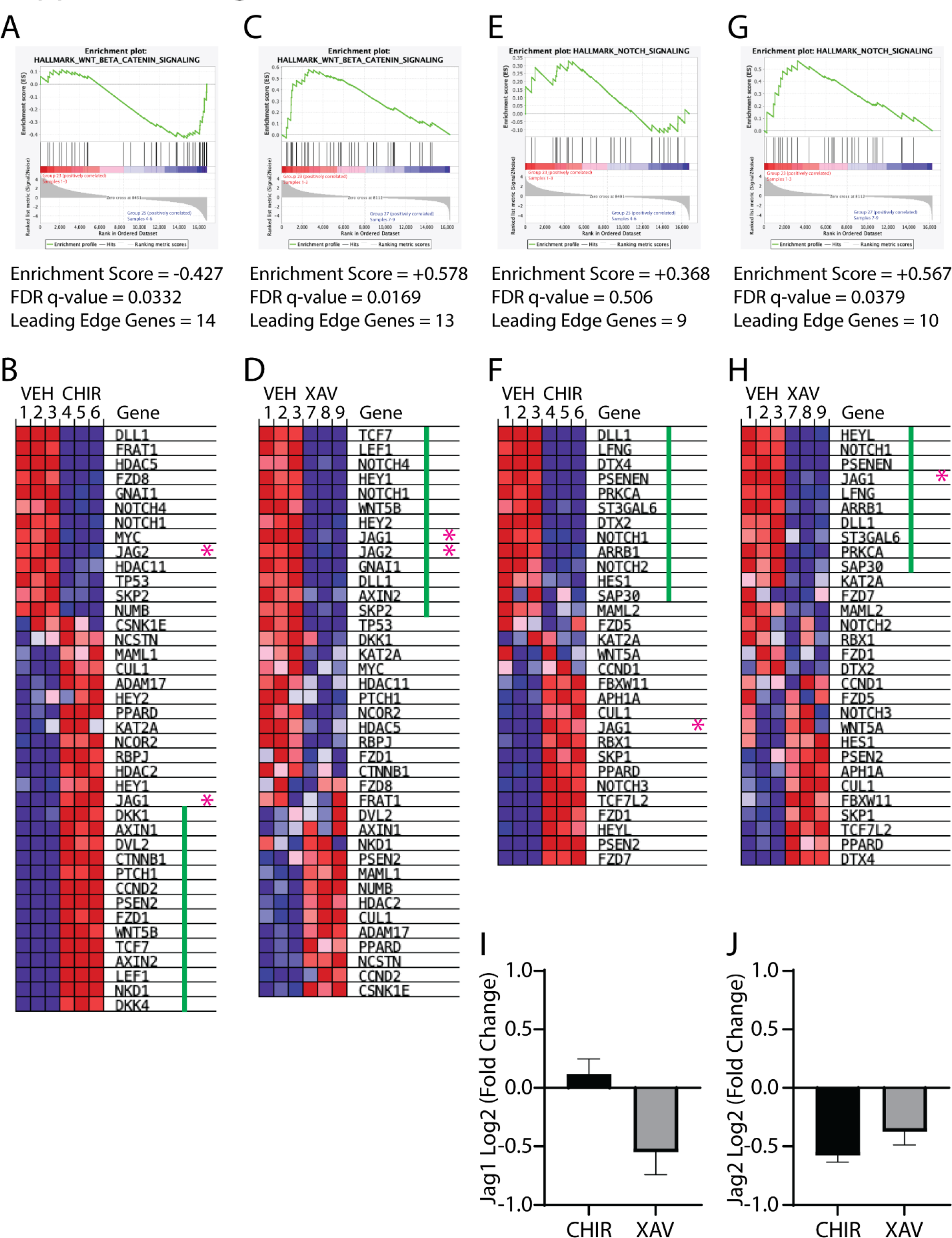
Gene expression and gene set enrichment analysis. ALI cultures were treated with vehicle, 10 µM CHIR or 10 µM XAV on differentiation days 4 and 6 and RNA was purified on day 8. Transcriptional changes were interrogated by RNA-sequencing and Gene Set Expression Analysis (GSEA). **A.** GSEA for WNT/β-catenin genes in vehicle and CHIR treated cultures. **B.**Heat map representation of WNT/β-catenin genes in vehicle and CHIR treated cultures. Each column is a sample: samples 1-3 vehicle treated (VEH), samples 4-6 CHIR treated. Each row is a gene. Red indicates upregulation. Blue indicates downregulation. Leading-edge genes are indicated by the green line. Jag1 and Jag2 are indicated by the asterisks. **C.** GSEA for WNT/β-catenin genes in vehicle and XAV treated cultures. **D.**Heat map representation of WNT/β-catenin genes in vehicle and XAV treated cultures. Each column is a sample: samples 1-3 vehicle (VEH treated), samples 7-9 XAV treated. Each row is a gene. Red indicates upregulation. Blue indicates downregulation. Leading-edge genes are indicated by the green line. Jag1 and Jag2 are indicated by the asterisks. **E.**GSEA for Notch genes in vehicle and CHIR treated cultures. **F.** Heat map representation of Notch genes in vehicle and CHIR treated cultures. Each column is a sample: samples 1-3 vehicle (VEH treated), samples 4-6 CHIR treated. Each row is a gene. Red indicates upregulation. Blue indicates downregulation. Leading-edge genes are indicated by the green line. Jag1 and Jag2 are indicated by the asterisks. **G.** GSEA for Notch genes in vehicle and XAV treated cultures. **H.** Heat map representation of Notch genes in vehicle and XAV treated cultures. Each column is a sample: samples 1-3 vehicle (VEH treated), samples 7-9 XAV treated. Each row is a gene. Red indicates upregulation. Blue indicates downregulation. Leading-edge genes are indicated by the green line. Jag1 and Jag2 are indicated by the asterisks. **I.** Analysis of *Jag1* gene expression in vehicle, CHIR, and XAV treated cultures. Data are presented as the Log2(fold change). Mean ± SD (n=3). No significant differences were detected. **J.** Analysis of *Jag2* gene expression in vehicle, CHIR, and XAV treated cultures. Data are presented as the Log2(fold change). Mean ± SD (n=3). No significant differences were detected.

**Supplemental Figure 6:**
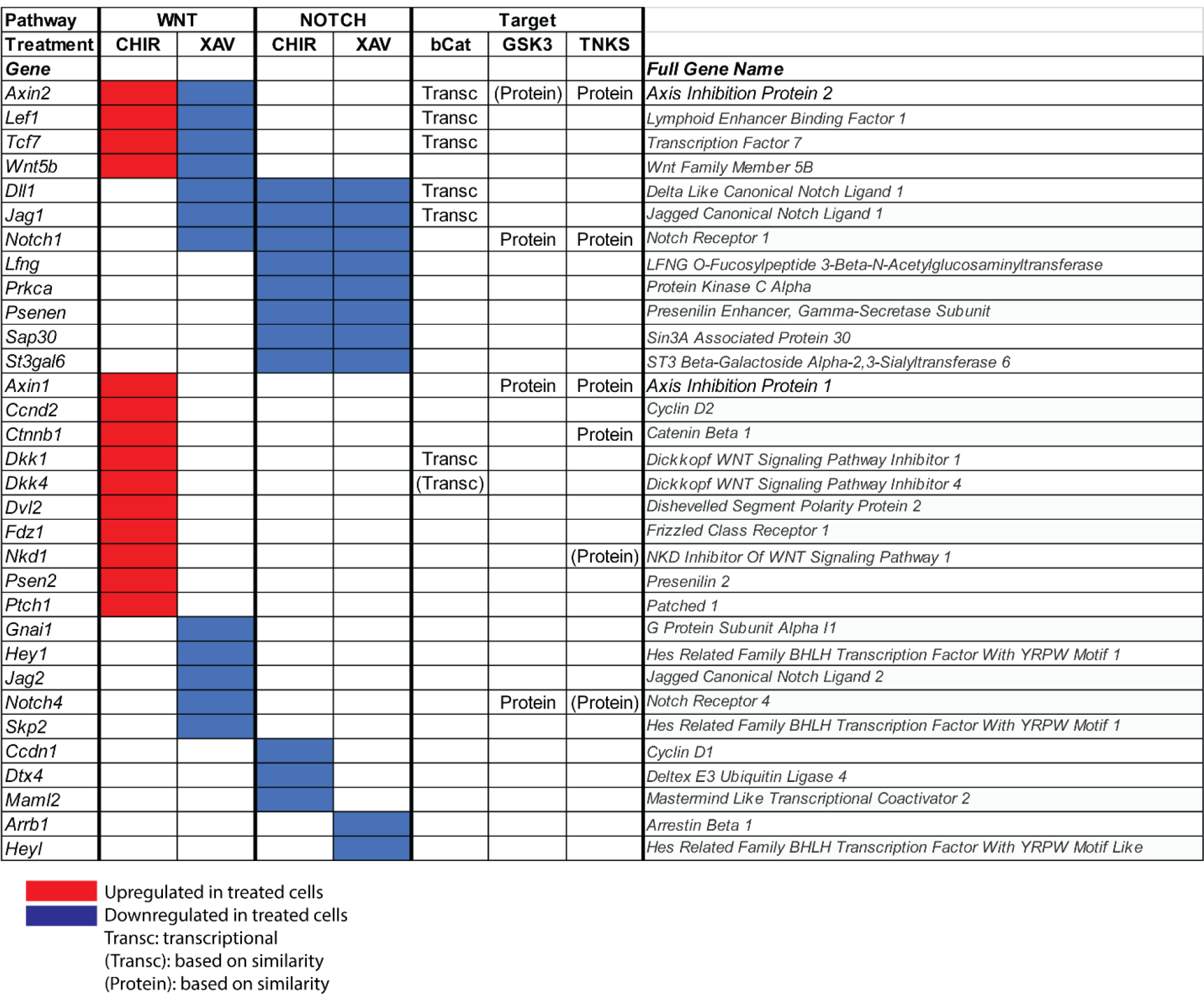
Analysis of leading-edge genes. ALI cultures were treated with vehicle, 10 µM CHIR or 10 µM XAV on differentiation days 4 and 6 and RNA was purified on day 8. Transcriptional changes were interrogated by RNA-sequencing and Gene Set Expression Analysis (GSEA). Leading-edge genes, which determine the gene set Enrichment Score, were extracted and organized to indicate similarities/differences across the treatments and gene sets. Red indicates downregulation. Blue indicates upregulation. Transc: Transcriptional targets of WNT/β-catenin were based on literature reports. (transc): WNT/β-catenin targets identified by similarity. Protein: Targets of glycogen synthase kinase (GSK3) and tankyrase (TNKS) were based on literature reports. (protein): GSK3 and TNKS targets identified by similarity.

**Supplemental Figure 7:**
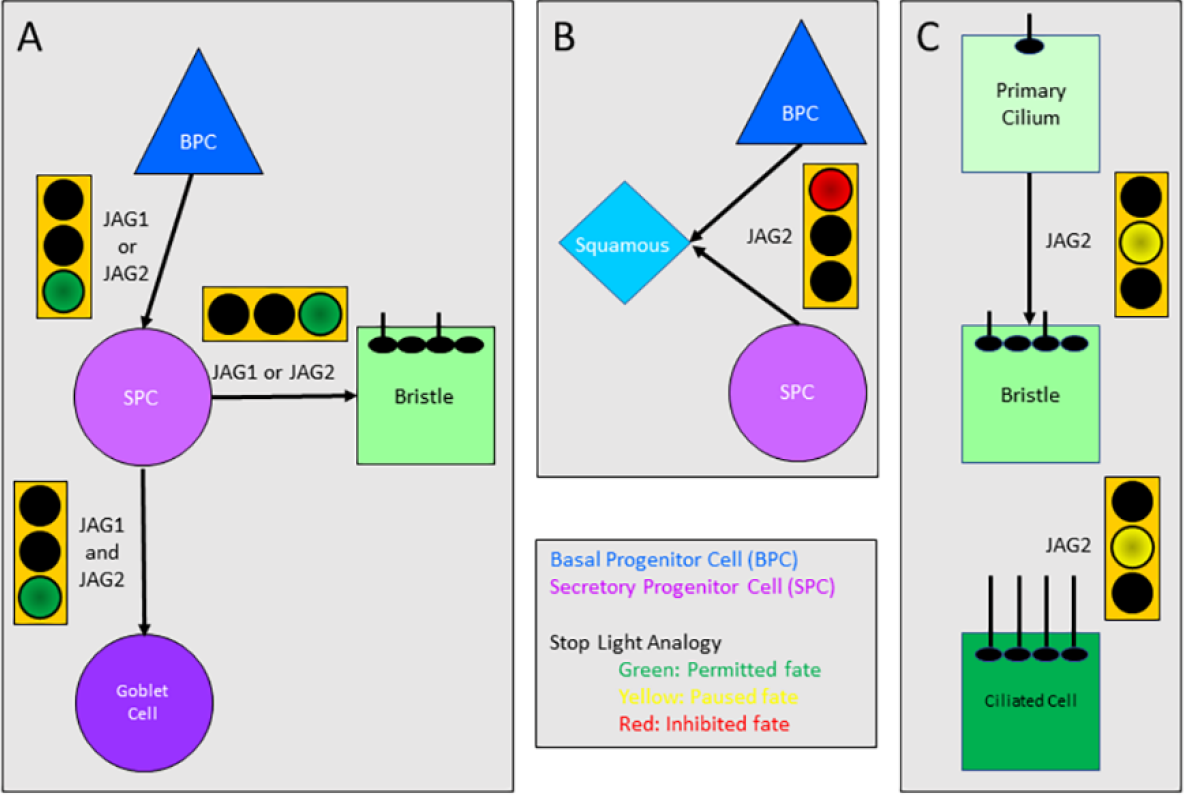
Roles for JAG1 and JAG2 in tracheobronchial cell fate decisions. **A**. JAG1 or JAG2 permit Basal Progenitor Cell (BPC) differentiation to a Secretory Progenitor Cell (SPC) and SPC differentiation to a bristle cell (green light). Both JAG1 and JAG2 are required for SPC differentiation to a goblet cell (green light). **B**. JAG2 inhibits the Squamous Cell fate (red light). **C.** JAG2 pauses progression from one ciliated cell differentiation intermediate to the next (yellow light).

